# Microbial odours activate protective immune response in *Caenorhabditis elegans* via specific olfactory neurons

**DOI:** 10.64898/2026.07.20.739438

**Authors:** Siddharth R. Venkatesh, Varsha Singh

## Abstract

Caenorhabditis elegans, like other animals, relies on attractive odours for foraging and on aversive odours for avoidance of pathogens. An odour produced by pathogenic bacterium Pseudomonoas aeruginosa induces immune response as well as avoidance in C. elegans via AWB olfactory neurons. We asked whether AWB neurons provide broader immunity to a wide range of pathogens. We activated AWB neurons using three chemically distinct bacterial odours and found that activation indeed induces protective immunity. Conversely, silencing of AWB neurons early during infection enhances susceptibility to infections. Mechanistic investigation revealed that the activation of AWB using odours upregulates detoxification pathways and other protective pathways in non-neuronal tissues. Specifically, odour exposure activates UDP-glucuronosyltransferase UGT-18 in the intestine that protects worms from the phenazine toxin of Pseudomonas aeruginosa and promotes broader immunity to Gram-negative bacteria (P. aeruginosa and Salmonella enterica), Gram-positive bacteria, (Enterococcus faecalis and Staphylococcus aureus), and yeast (Cryptococcus neoformans). Altogether, our findings highlights microbial odours as non-canonical molecular patterns for activating immune responses.

## INTRODUCTION

Pattern recognition during infection is the cornerstone of immune defence, enabling the timely initiation of innate protective responses. This recognition typically involves the detection of pathogen-associated molecular patterns (PAMPs) by pattern recognition receptors (PRRs)^1–4^. Invertebrates, *Drosophila melanogaster* and *Caenorhabditis elegans*^5,6^, have been instrumental in uncovering both canonical and non-canonical mechanisms of pathogen detection,. In vertebrates, a diverse array of PRRs, including Toll-like receptors (TLRs), RIG-I-like receptors (RLRs), absent in melanoma 2 (AIM2)-like receptors (ALRs), and nucleotide oligomerisation domain (Nod)-like receptors (NLRs), mediate the detection of a broad spectrum of PAMPs^1,7^. While many of these PRRs are conserved in *D. melanogaster*^6,8^, they are largely absent in *C. elegans*^9^. The PRRS detect a diverse range of molecules including lipopolysaccharide, DNA and RNA and peptides. Recently smaller molecules such as odours have been shown to elicit immune response in *C. elegans*. It is unclear if other odors from pathogenic microbes are PAMPs and whether this mechanism is widespeard in the animal kingdom.

*C. elegans*, a free-living, bacterivorous nematode, is found in decomposing vegetation in and around soils. Like other invertebrates, *C. elegans* relies exclusively on innate and inducible immunity to counter pathogenic microbes in its habitat. Its defence strategies include both a flight and a fight response. The former is an aversive response to specific pathogens, while the latter involves the induction of antimicrobial effectors such as lysozymes, lectins, antimicrobial peptides (AMPs), and cytoprotective molecules such as superoxide dismutase and cytochrome P450. Notably, *C. elegans* has few canonical PRRs^9^. The sole Toll-like receptor homologue, TOL-1, is dispensable for defence against most pathogens, except for *Salmonella enterica*^10^ and *Serratia marcescens*^11,12^. Meanwhile, the RIG-I-like receptor DRH-1 detects viral replication intermediates to activate an intracellular response^13^. C-type lectin-like domain (CTLD) proteins, such as CLEC-39 and CLEC-49, contribute to immune defence against *S. marcescens* by binding to live bacteria^14^. Despite these insights, the molecular underpinnings of pathogen recognition and ensuing activation of specific or broader immunity in *C. elegans* remain poorly understood. Recent evidence suggests the roles of non-canonical pathways for pathogens recognition and immune modulation, particularly through G protein-coupled receptors (GPCRs)^15,16^ in amphid sensory neurons^17,18^. Notably, a previous study demonstrated that an 1-undecene odour produced by *Pseudomonas aeruginosa*, is a molecular PAMP for *C. elegans*^19^. This prompted us to consider if odours from other pathogens act as molecular patterns for GPCRs in olfactory neurons.

Olfaction plays a pivotal role in the life history of *C. elegans*, guiding behaviours such as foraging^20^ and social behaviour^21^ as well as life span. The amphid sensory organ harbours three pairs of olfactory neurons, AWA, AWB, and AWC, expressing overlapping but distinct sets of GPCRs. While AWA and AWC predominantly detect attractive odours, AWB neurons specialise in sensing aversive odours and mediating pathogen avoidance. Notably, AWB neurons are critical not only for initiating avoidance behaviours but also for activating immune responses against *P*. *aeruginosa* by detecting 1-undecene odour produced uniquely by this pathogen^19^. Priming by 1-undecene induces induction of infection regulated genes (irg) in manner edpendent on AWB neurons. These findings raise the possibility that other AWB neuron-sensed odour are PAMPs necessary for innate immune activation.

In this study, we asked whether activation of AWB olfactory neurons by pathogen-derived aversive odours could prime innate immunity against a diverse range of pathogens. Using using two distinct odours for the activation of AWB neurons, we could observe protective immunity against 5 different pathogens. By blocking AWB activation during infection, we established temporal requirement of odour primed immunity during the early phases of host defence. Analyses of whole animal transcriptome during odour-mediated priming event revealed significant upregulation of immune genes, particularly those associated with xenobiotic stress responses and detoxification. In particular, we found that a UDP-glucuronosyltransferase UGT- 18, involved in detoxification, is activated via the AWB olfactory neurons. UGT-18 is expressed in the intestine and helps in protection from specific toxins including phenazine produced by P. aeruginosa. Altogether, our findings highlight a novel role for microbial odours and olfactory neurons in coordinating detoxification and immune pathways in response to pathogens.

## RESULTS

### Activating AWB olfactory neurons by microbial odours promotes survival of *C. elegans* during infection by broad range of pathogens

1-Undecene (Und), a *P*. *aeruginosa*-specific odour, is a pathogen-specific molecular pattern working via AWB olfactory neurons of *C. elegans*^19^. Pathogenci bacteria also produce odours. We hypothesized that one of the functions of AWB neurons, responsible for sening aversive odours, is to prime broader protective immunity to microbial pathogens. Specifically, we asked whether activation of AWB by diverse odours can induce immunity to broad classes of pathogens. We exposed worms to Und (an alkene) and 2-nonanone or Non (a ketone) for 2 hours and then assessed the survival of worms on subsequent infection with 5 different pathogens individually. First, we utililized Und to activate AWB neurons (**Figure 1A**), followed by infection with *P*. *aeruginosa*. Und-exposed worms were significantly more resistant to *P*. *aeruginosa* infections compared to naïve (unexposed) worms, as shown earlier (**Figure 1B**)^19^. However, odour-induced immunity was not observed when worms lacking AWB neurons (AWB(-) were exposed to Und (**Figure 1C**), confirming the requirement of the AWB neurons. Pre-exposure to Und also significantly improved the survival of worms on another Gram-negative pathogen, *Salmonella enterica*, two Gram-positive pathogens, *Enterococcus faecalis* and *Staphylococcus aureus,* and pathogenic yeast *Cryptococcus neoformans* (**Figures 1D-G**). This indicated that the activation of immune pathways by Und odour activates broader immunity. To confirm this, we used another AWB-sensed bacterial odour, 2-nonanone, and assayed survival during subsequent infection. We found that 2- nonanone exposed worms also displayed better survival on all 5 pathogens tested compared to naïve worms (**Figures S1A-E, also see Table S1**). These experiments indicated that activating AWB neurons via two different odour ligands is sufficient to enhance survival from infections with pathogens.

**Figure 1.**
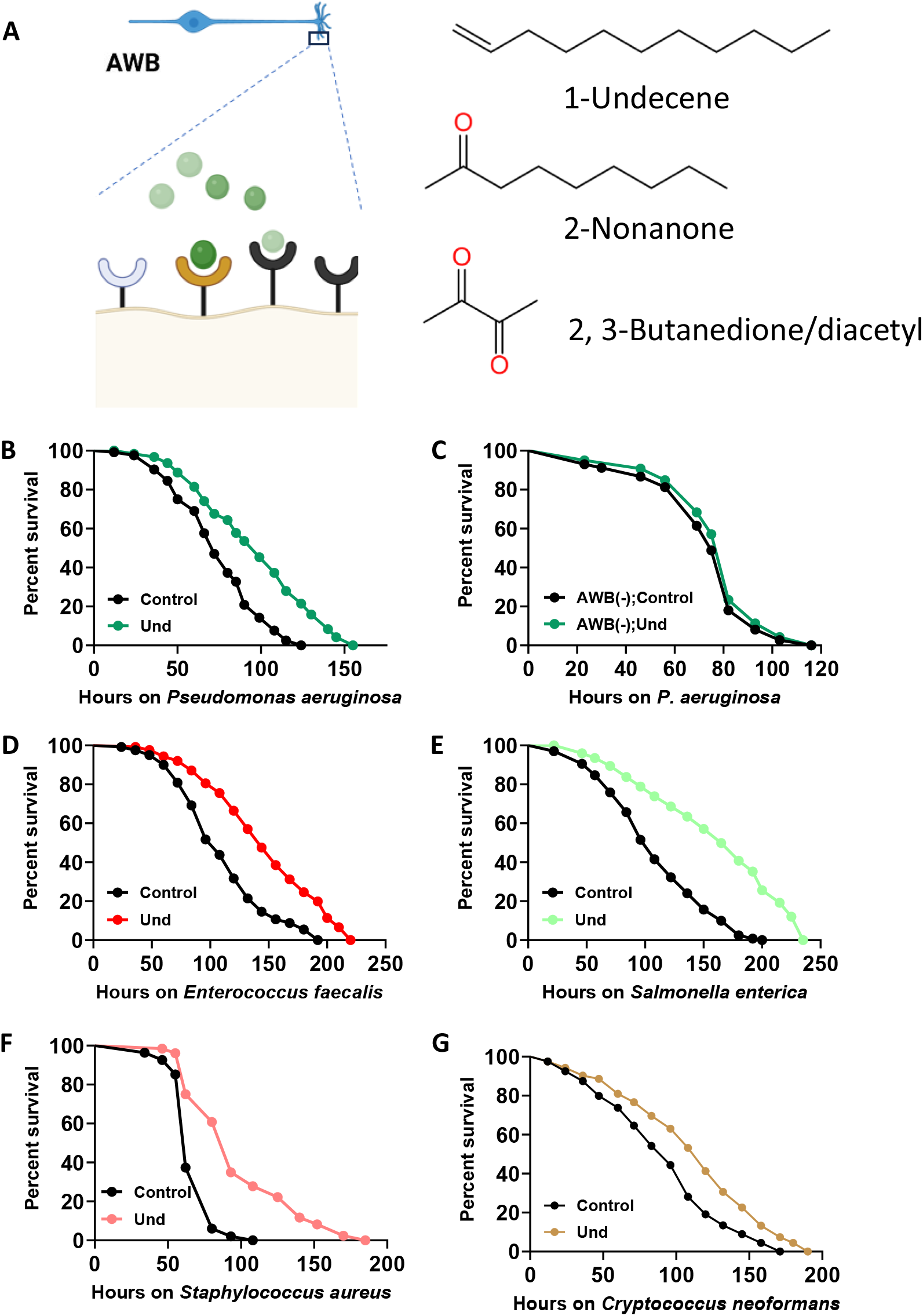
Activating AWB neurons using pathogen-derived odours protects worms during subsequent infection. (A) Schematic of AWB activation using odours. (B) Kapler-Meier survival curves of control and 1-undecene (Und) exposed (B) WT (wild-type; *P* < 0.0001) and (C) AWB(-) (*P* = 0.1712) worms on *Pseudomonas aeruginosa*. Kapler-Meier survival curves of control and 1-undecene exposed WT worms on (D) *Enterococcus faecalis* (*P* < 0.0001), (E) *Salmonella enterica (P* < 0.0001), (F) *Staphylococcus aureus* (*P* < 0.0001), and (G) *Cryptococcus neoformans* (*P = 0.0002)*.

Next, we sought to confirm that the immunity-boosting role of AWB neurons by genetically programming it (AWB) to sense an odour normally sensed by AWA neurons. We utilized a GPCR misexpression line called CX3877 ((*odr-10(ky225)*; kyIs156 [*str-1p*::*odr- 10*(cDNA)::GFP]). Wild-type (WT) worms are attracted to attractive odour diacetyl (DA) via olfactory receptor ODR-10 expressed in the AWA neurons^22^. CX3877 strain has a mutated copy of *odr-10* in the genome, and wild type of of *odr-10* is (mis)expressed only in the AWB neurons^22^ (**Figure 2A**). As a result, DA causes an aversion response in CX3877 worms^22^. We found that exposing WT worms to DA prior to infection does not alter their survival on pathogens (**Figures 2B-E**), suggesting that AWA neurons do not regulate immunity pathways in these worms. However, pre-exposure of CX3877 strain to DA significantly enhanced their survival on pathogens compared to naïve CX3877 worms (**Figures 2B-E**). This demonstrated that the activation of AWB neurons (Und, Non or DA) by various means is sufficient to activate broader immunity against infections.

**Figure 2.**
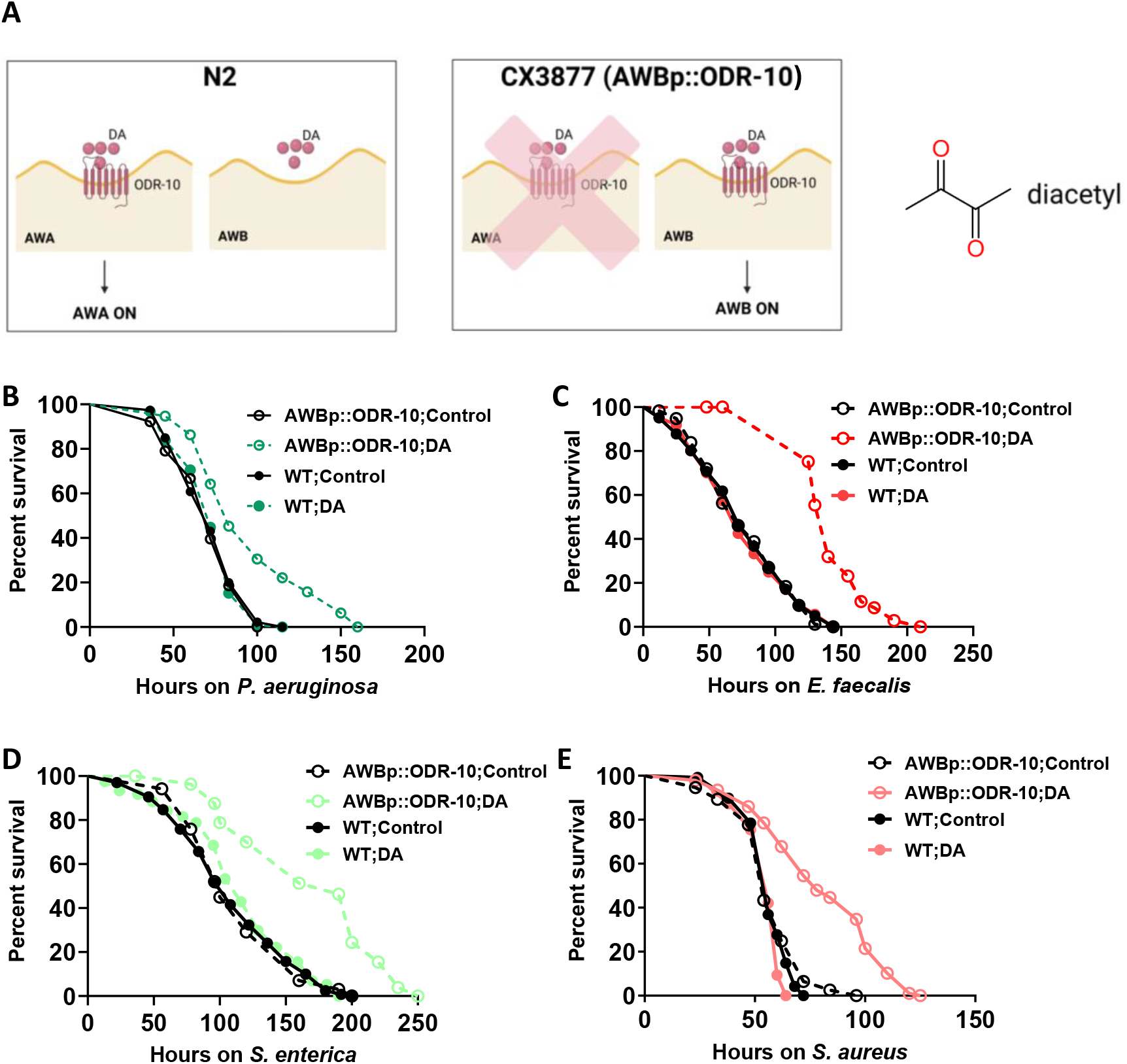
Odour activated protective immunity is AWB neuron-dependent. (A) Schematics of AWB activation using diacetyl (DA) in WT (wild-type, N2) and CX3877 (*odr-10(ky225)*; kyIs156 [*str-1p*::*odr-10*(cDNA)::GFP]) worms and the structure of diacetyl. Kapler-Meier survival curves of WT and CX3877 control and DA exposed worms on (A) *P. aeruginosa* (*P* < 0.0001), (B) *E. faecalis* (*P* < 0.0001), (C) *S. enterica* (*P* < 0.0001), and (D) *S. aureus* (*P* < 0.0001).

### Olfactory neuron function is needed early during infection for protective immunity

Conventional PAMPs activate innate immune pathways early during infection. To test if volatile PAMPs work similarly, we resorted to silencing AWB neurons at different times during infection. By expressing a histamine-gated chloride channel (HisCl) in AWB neurons specifically, we could silence (hyperpolarise) these neurons by applying histamine (HA) for discrete periods of time (**Figure 3A**)^23^. We created a strain exclusively expressing *Drosophila melanogaster* histamine-gated chloride channel 1 (HisCl1) in the AWB neurons (AWBp::HisCl1). We confirmed the effectiveness of silencing by studying the response of HA-treated and untreated VSL2301 (AWBp::HisCl1) worms to Und and Non. While control worms were strongly repulsed by either Und and Non, worms treated with histamine for 4 h or 8h had muted repulsion to these odours (**Figures 3B and 3C**) indicating effective silencing of AWB neurons. However, histamine treated worms displayed WT-like chemotaxis to an AWA-sensed odour, DA (**Figure 3D**), suggesting that silencing was restricted to AWB neurons.

**Figure 3.**
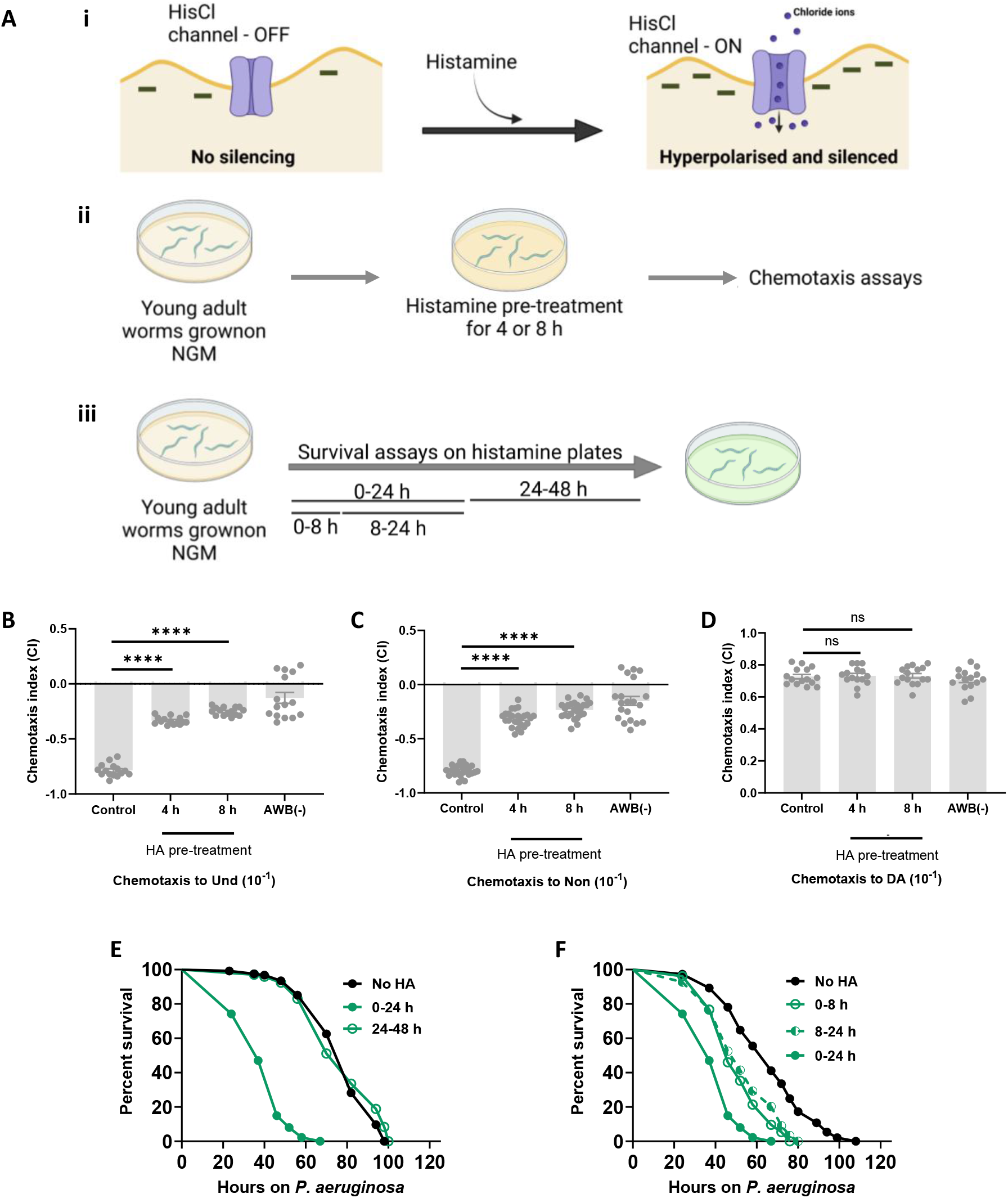
Silencing of AWB neurons early during infection reduces survival on pathogens. (A) Schematics of Histamine-gated chloride channel (HisCl)-mediated silencing of neurons. (B-D) Bar graphs depicting the chemotaxis index of VSL2301 (AWBp::HisCl, histamine responsive AWB neurons) with or without histamine (HA) and AWB(-) worms for (B) 1-undecene (Und), (C) 2-nonanone (Non) and (D) diacetyl (DA). The error bars represent the standard error of the mean (SEM). Statistical analysis was performed using one-way ANOVA followed by a post-hoc Dunnet test. \*\*\*\**P* < 0.0001; ns, not significant. Kaplan-Meier survival curves of VSL2301 worms or VSL2301 worms treated with HA for (C) 0-24 (*P* < 0.0001) and 24-48 h (*P* = 0.4994) and (D) 0-8 (*P* < 0.0001), 8-24 (*P* < 0.0001), and 0-24 h (*P* < 0.0001). Same set of 0-24 hours HA treated worms are shown in panels E and F.

To identify the temporal window of requirement for AWB neuronal function during infection, we silenced AWB neurons using histamine for 0-24 hours or 24-48 hours during infection with P*. aeruginosa* and scored survival. Interestingly, VSL2301 worms treated with histamine for 0-24 hours during infection with *P*. *aeruginosa* were significantly more susceptible to infection than untreated VSL2301 worms (**Figure 3E**). However, worms treated with histamine during 24-48 hours of infection had survival comparable to untreated VSL2301 worms, suggesting that AWB neuronal function was relevant in the first 24 hours. To further demarcate the exact window within which AWB function is required for protective immunity, we silenced AWB neurons for 0-8 or 8-24 hours during infection. Both these treatment regimens rendered the worms more susceptible than control worms but less so than worms treated for 0-24 hours (**Figure 3F**). These experiments suggest that AWB functions within the first 24 hours of infection to protect worms against pathogens.

### Odour-primed immune regulon in *C. elegans:* Pathogen odours activate detoxification pathways and immune-response genes via olfactory neurons

To identify molecular players of AWB-mediated protective immunity, we analysed the bulk transcriptome of WT worms exposed to Und or Non, for just 90 minutes to prime immune response, and compared them to the transcriptome of naïve WT worms. Principal component analysis (PCA) revealed that the transcriptome of the Und and Non exposed groups clustered along PC1, which accounted for nearly 24% of the variation within all datasets (**Figure 4A**). We constructed a volcano plot using the SRplot tool to analyse the differentially expressed genes (DEGs)^24^. **Figures 4B and D** show that 1054 and 1374 genes were significantly upregulated, while 1493 and 1498 genes were significantly downregulated in the Non and Und groups, respectively, compared to Con worms. Gene ontology (GO) analyses indicated that biological processes, such as “Response to heat,” “Response to stress,” “Response to Gram-positive bacterium,” “Response to Gram-negative bacterium,” and “Innate immune response,” were significantly enriched in Und worms (**Figure 4C**). Similarly, GO biological processes, such as “Defence response,” “Response to Gram-positive bacterium,” “Response to Gram-negative bacterium,” “Innate immune response,” and “Heat response,” were enriched in the Non worms (**Figure 4D**). Notably, innate immune genes included those encoding proteins involved in detoxification, toxin resistance, and xenobiotic stress responses (Supplementary Table S2; WormExp analysis). This analysis indicated that a large subset of genes with known and potential involvement in stress sresponse and immune response are activated during brief priming with odours.

**Figure 4.**
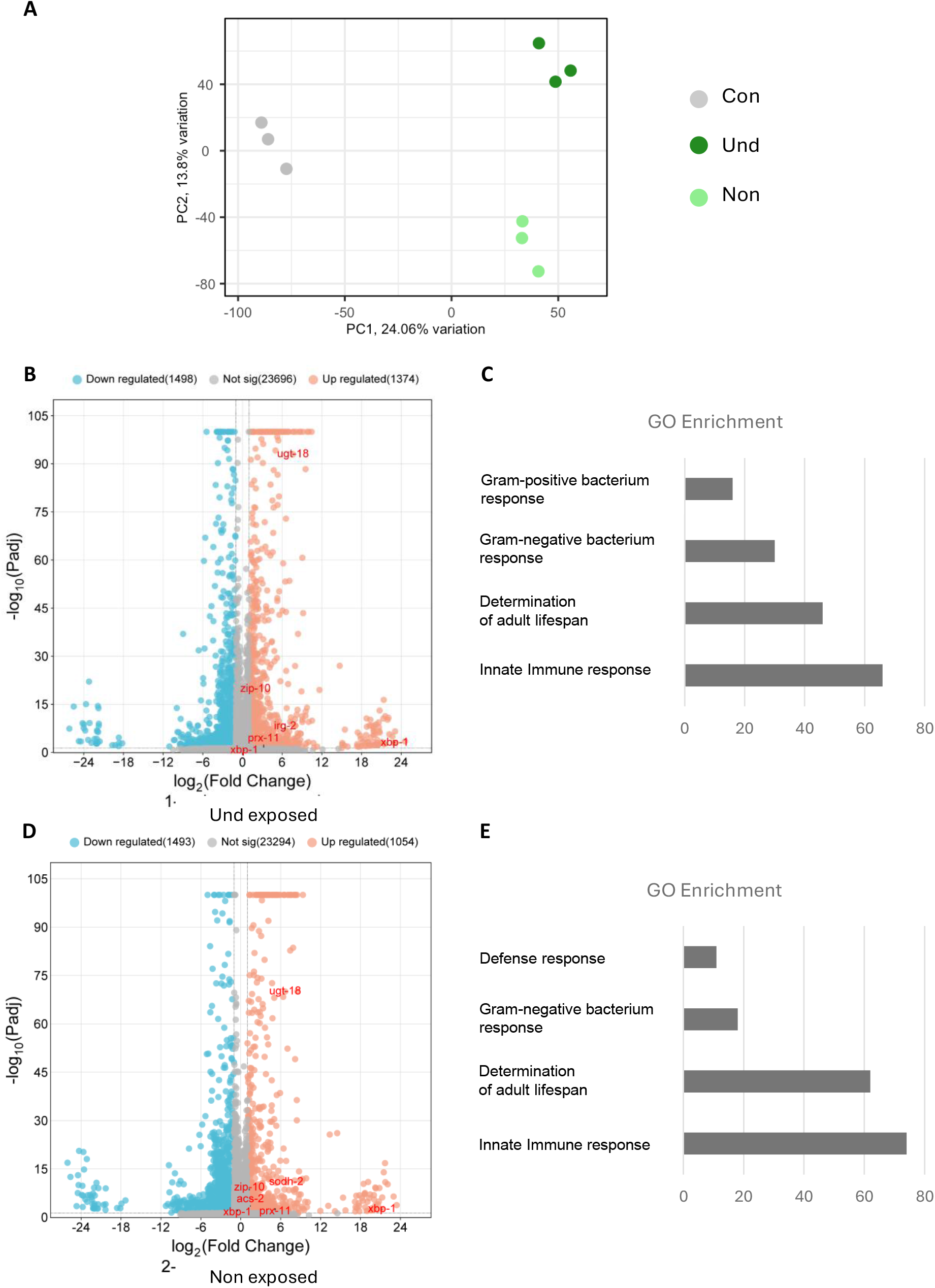
Immune-related genes are enriched in volatile-exposed worms. (A) Principal component analysis of worms exposed to 1-undecene (Und) and 2-nonanone (Non) and unexposed (Con) worms. (B) Volcano plot of Und exposed worms and the corresponding (C) enriched gene ontology (GO) terms for upregulated genes. (D) Volcano plot of Non exposed worms and the corresponding (E) enriched GO terms for upregulated genes.

To identify specific immune response genes from amongst odour primed transcripts, we compared the transcriptome of Und and Non exposed worms with the transcriptome of worms infected with three broad classes of pathogens^25^, Gram-ve bacterium *P*. *aeruginosa*, Gram +ve bacterium *E*. *faecalis*, and pathogenic yeast *C*. *neoformans*. The Venn diagram in **Figure 5A** depicts the number of genes upregulated under these conditions. We identified 33 genes that were upregulated during 8 hour infections with either of the pathogens, as well as by priming with odours for 90 minutes. A majority of these genes are categorised as “immunity-related” genes by GO analysis. We validated the upregulation of 17 of these genes by quantitative real-time PCR (qRT-PCR) in worms exposed to Und and Non (**Figures 5B and S2**). Some of these candidate genes encoded detoxification proteins, such as cytochrome P450 (CYP-34A9, CYP- 34A10, etc.), UDP-glucuronosyltransferase (UGT-18), alcohol dehydrogenase (SODH-2), and cycteine proteases (CPR-3). To confirm that these transcripts were upregulated due to the activation of AWB neurons, we also performed qRT-PCR in odour exposed AWB(-) worms. As shown in **Figures 5B**, AWB(-) worms did not show significant upregulation of these genes upon Und or Non exposure. We created a fluorescence transcriptional reporter for the most upregulated gene, *ugt-18* for further anaysis. A GFP reporter *ugt-18* (VSL2401), showed that both 1-undecene and -nonanone odours lead to 4-10 fold upregulation (**Figure S3, A-B**). Taken together, the transcriptome analyses indicated that Und, as well as Non, prime immune response comprised of many cytoprotective pathways in *C. elegans*.

**Figure 5.**
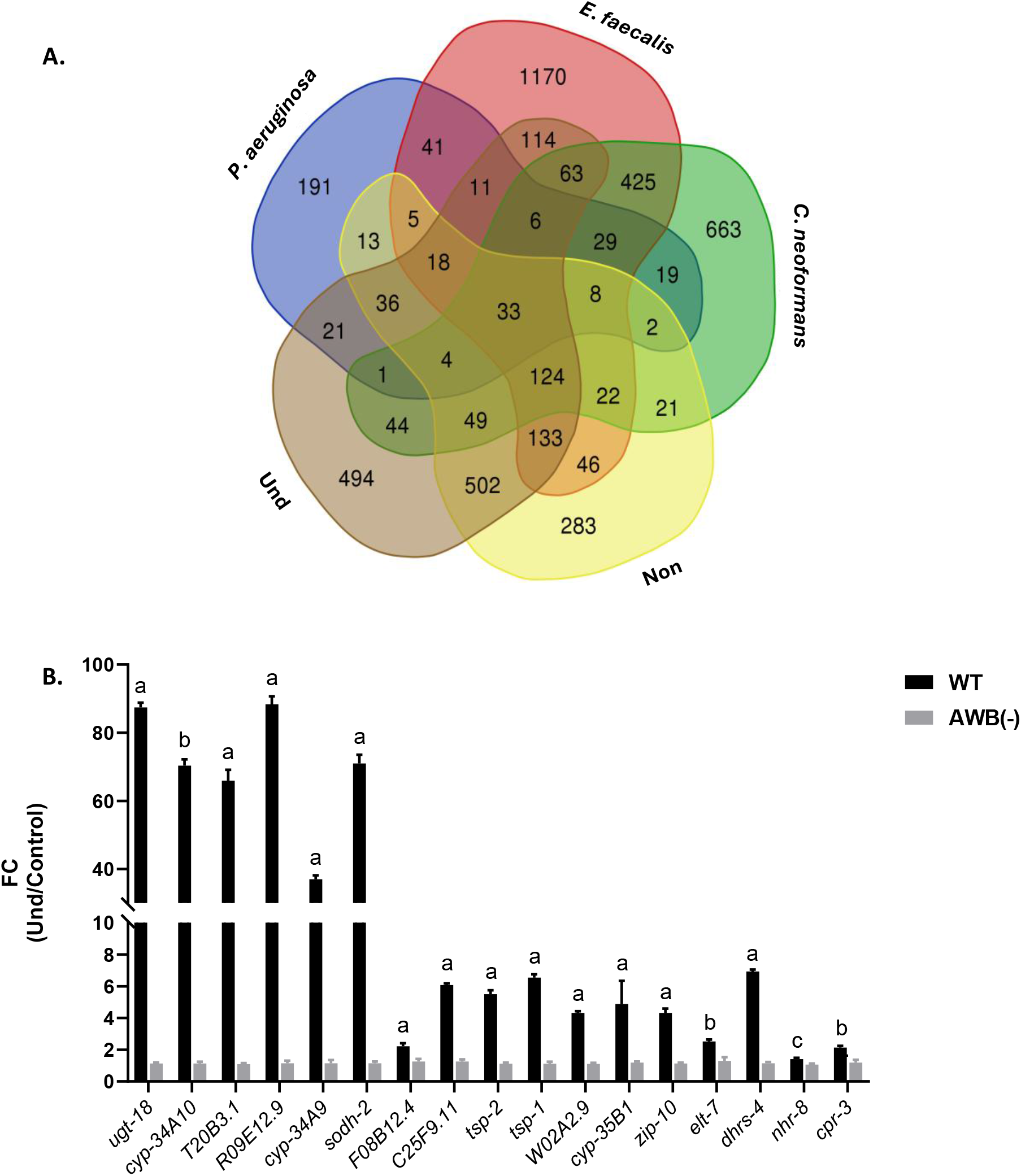
Odour-primed immune regulon in *C. elegans*. (A) Venn diagram for the genes upregulated in worms infected with *P. aeruginosa*, *E. faecalis*, and *C. neoformans* and worms exposed to 1-undecene (Und) and 2-nonanone (Non). (B) qRT-PCR validation of select immune-response genes and transcription factors of WT (wild-type) and AWB(-) worms exposed to 1-undecene (Und). Statistics were calculated using an unpaired Student’s t-test. a, *P* < 0.0001; b, *P* < 0.001; c, *P* < 0.01; d, *P* < 0.05. The error bars indicate the standard error of the mean (SEM).

### AWB-regulated gene encoding UDP-glucuronosyltransferase UGT-18 protects worms from phenazine toxin of *P*. *aeruginosa*

To establish the relevance of odour primed immune regulon in survival during infection, we tested the role of 9 genes in survival on *P. aeruginosa*. RNAi-mediated inhibition of *ugt-18*, *sodh-2*, *R09E12.9*, *T20B3.1*, *W02A2.9*, and *cpr-3* significantly reduced survival of worms on *P*. *aeruginosa* (**Figures 6A-F**), with the maximum impact observed with *ugt-18* RNAi. However, the RNAi of *tsp-1*, F08B12.4, and C25F9.11 did not affect the survival of worms on *P*. *aeruginosa* (**Figure 6G-I**). This indicated 2/3 of odour primed genes had a role in survival during infection with UDP-glucuronosyltransferase *ugt-18* being the most effective against *P. aeruginosa*.

**Figure 6.**
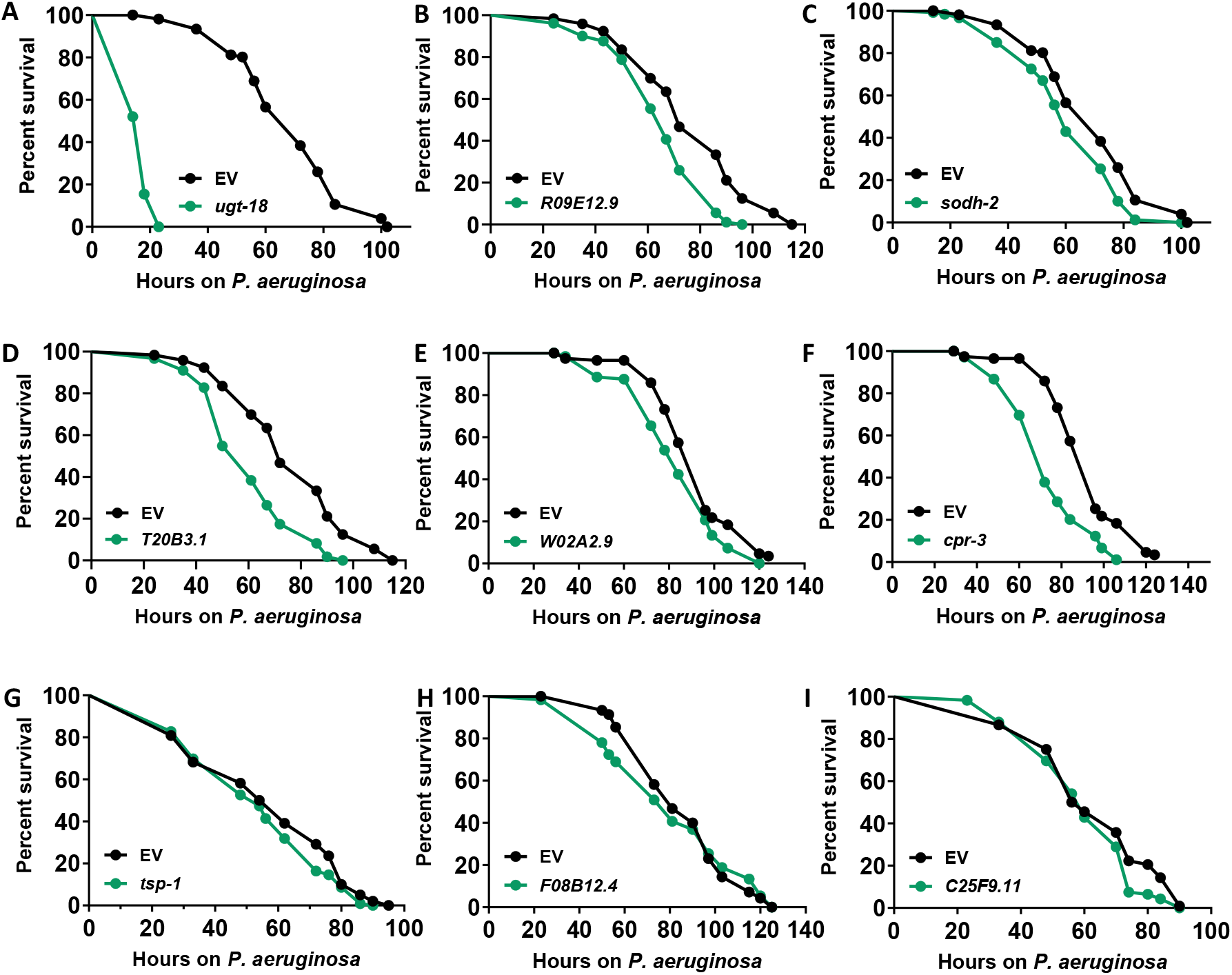
AWB-regulated genes are required for survival of C. elegans from infection by *Pseudomonas aeruginosa*. Kaplan-Meier survival curves of (A) *ugt-18* (*P* < 0.0001), (B) *R09E12.9* (*P* < 0.0001), (C) *sodh-2* (*P* = 0.0015), (D) *T20B3.1* (*P* < 0.0001), (E) *W02A2.9* (*P* = 0.0030), (F) *cpr-3* (*P* < 0.0001), (G) *tsp-1* (*P* = 0.1348), (H) *F08B12.4* (*P* = 0.4263), and (I) *C25F9.11* (*P* = 0.614) knockdown worms on *Pseudomonas aeruginosa*.

UDP-glucuronosyltransferases (UGTs) are a family of enzymes that play a crucial role in the detoxification of endogenous and exogenous molecules. UGTs catalyse the transfer of glucuronic acid from the co-substrate UDP-glucuronic acid to various molecules, including drugs, xenobiotics, and toxins^26^. Glucuronidation process enhances the water solubility of lipophilic substances, facilitating their subsequent excretion or breakdown and detoxification^27^. In *C*. *elegans*, some of the UGTs are important for the detoxification of toxins and drugs^28,29^, including phenazine, a secondary metabolite of *P*. *aeruginosa* essential for its virulence^30–32^ and cytotoxicity of host cells. Therefore, we tested whether UGT-18 protects worms from direct exposure to phenazine.

We set up a chronic phenazine exposure assay ofr C. elegans. We found a time-dependent toxicity of phenazines such that 33% of WT worms were dead at 9 hours of exposure to 180 μM phenazine while control worms were alive (**Figure 7A-B**). We used this time point in subsequent assays. We found that *ugt-18* RNAi worms had significantly reduced survival on phenazine at 9 hours than control worms (**Figure 7D-E**). We also observed a similar reduction in survival in the AWB(-) worms upon exposure to phenazine for 9 hours compared to WT worms (**Figure 7D and 7F**). Notably, Und-primed WT worms were significantly better protected against phenazine than unprimed WT worms (**Figure 7G-H**). Moreover, Und- mediated protection against phenazine was not observed in AWB(-) worms (**Figure 7H**), consistent with the role of AWB neurons-in the upregulation of protective pathways including *ugt-18*. To confirm the involvement of *ugt-18* in odour-primed worms, we studied the effect of Und exposure on survival of control and *ugt-18* RNAi worms upon phenazine exposure. We found that Und priming did not protect *ugt-18* RNAi against phenazine (**Figure 7G-I**). We made similar observations on the requirement of UGT-18 when AWB neurons were activated using Non odour followed by phenazine toxicity assay (**Figure S4A-B**). These assyas established the requirement of AWB neuron- intestinal UGT-18 axis as one of the major pathway for phenazine detoxification (**Figure 7J**).

**Figure 7.**
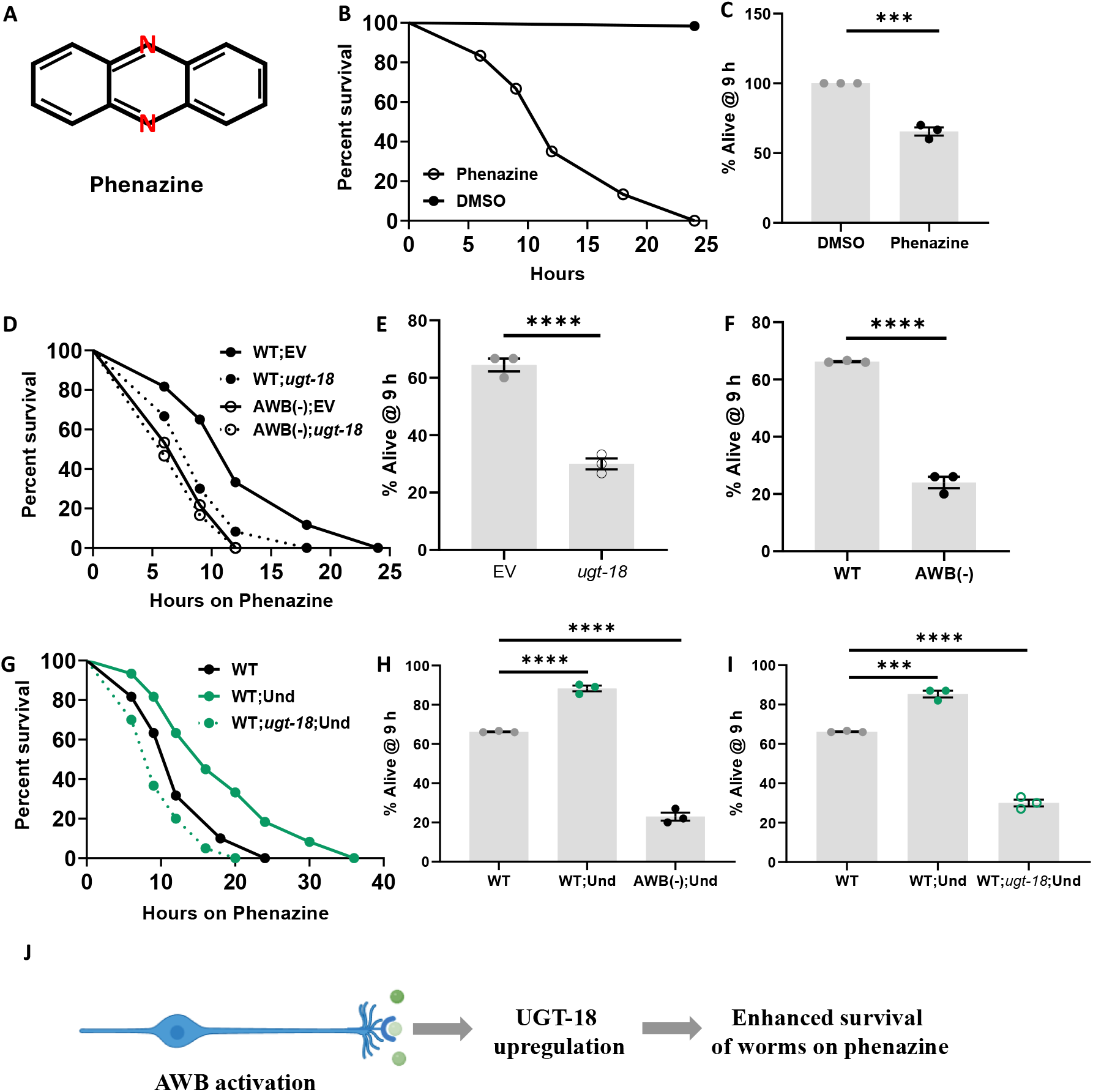
*ugt-18* is required for survival of worms from phenazine toxicity. (A) Structure of phenazine. (B) Kaplan-Meier survival curve of WT worms exposed to DMSO or phenazine (*P* < 0.0001). (C) Bar graph depicting the survival of WT worms on phenazine 9 h post exposure. (D) Kaplan-Meier survival curve of WT or AWB(-) worms with or without *ugt-18* knockdown (*P* < 0.0001) on phenazine. (E) Bar graph depicting the survival of WT worms with *ugt-18* RNAi knockdown on phenazine at 9 h. (F) Bar graph depicting the survival of WT and AWB(-) worms on phenazine at 9 h. Statistics for C, E, and F were performed using unpaired Student’s *t*-test with Welch’s correction. \*\*\**P* < 0.001; \*\*\*\**P* < 0.0001. (G) Kaplan-Meier survival curves for WT worms exposed to 1-undecene (Und) on phenazine, with or without *ugt-18* RNAi knockdown (*P* < 0.0001). (H) Bar graph depicting the survival of WT and AWB(-) worms on phenazine that were exposed to Und. (I) Bar graph depicting the survival of WT worms on phenazine, with or without *ugt-18* RNAi knockdown and exposure to Und. Statistics for H and I were performed using a one-way analysis of variance (ANOVA), followed by Dunnet test. \*\*\**P* < 0.001; \*\*\*\**P* < 0.0001; ns, not significant. The error bars indicate the standard error of the mean (SEM). (J) Schematic describing the upregulation of *ugt-18* upon AWB activation, which in turn promotes survival from phenazine toxicity.

### UGT-18 detoxification mechanism protects *C. elegans* against broad classes of pathogens

UGTs are enzymes with capability to detoxify diverse sets of molecules. We hypothesised that UGT-18 protects against Gram-negative and Gram-positive bacteria and pathogenic yeast by detoxifying diverse toxins they make. To test this hypothesis, we examined the survival phenotype of *ugt-18* RNAi worms on *E*. *faecalis*, *S*. *enterica*, *S*. *aureus,* and *C*. *neoformans.* WE found that *ugt-18* RNAi worms were significantly more susceptible than control worms on all pathogens (**Figures 8A-D**). Consistent with the requirement of UGT activity for protection against diverse pathogens, we observed significant upregulation in *ugt-18* mRNA levels in WT worms infected with *P*. *aeruginosa*, *E*. *faecalis*, *S*. *enterica*, *S*. *aureus*, and *C*. *neoformans* (**Figure 8E**), supporting the role for UGT-18 in during all microbial infections. Moreover, the induction in *ugt-18* mRNA levels was dependent on the presence of AWB olfactory neurons as AWB(-) worms exposed to these pathogens did not show any significant upregulation in *ugt-18* mRNA levels (**Figure 8E**). To confirm this we also examined VSL2401 (P*ugt-18*::GFP), and observed significant upregulation in GFP levels when exposed to all the tested pathogens (**Figures S5**). Altogether these experments indicated that UGT-18, a detoxifying enzymes, is upregulated in C. elegans intestine during infection and contributes to protective immunity against Gram negative bacteria, Gram positive bacteria and pathogenic yeast.

**Figure 8.**
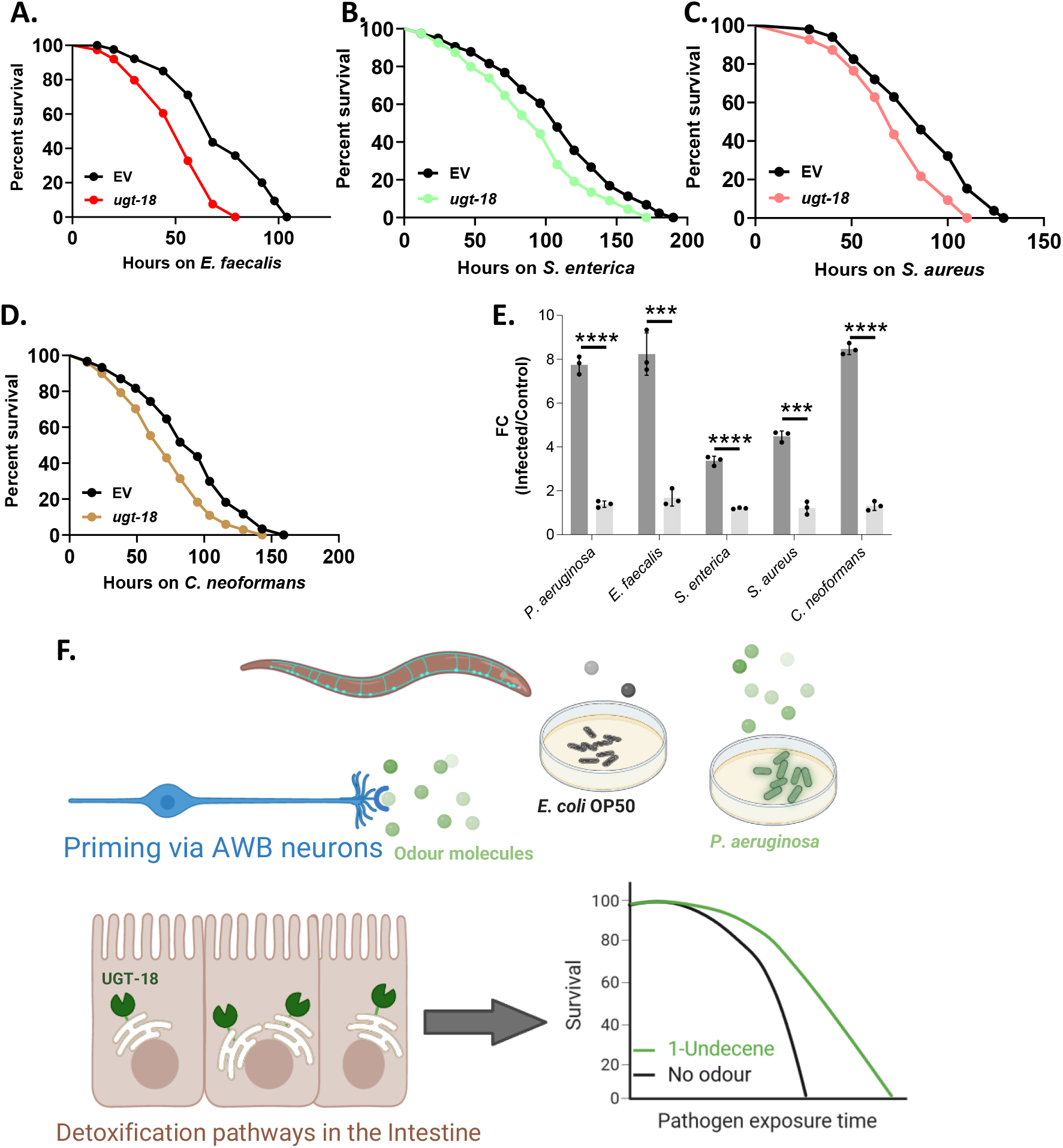
UGT-18 is required for survival on all tested pathogens. Kaplan-Meier survival curves of *ugt-18* knockdown animals on (A) *E. faecalis* (*P* < 0.0001), (B) *S. enterica* (*P* = 0.0026), (C) *S. aureus* (*P* = 0.0002), and (D) *C. neoformans* (*P* < 0.0001). (E) qRT-PCR data for *ugt-18* levels in WT (wild-type) worms exposed to *P. aeruginosa*, *E*. *faecalis*, *S*. *enterica*, *S*. *aureus*, and *C*. *neoformans*. (F) Working model for the induction of systemic protective pathways in C. elegans by priming of olfactory neurons by microbial odors. Activation of specific cytoprotective pathways such as uracil glycosyl transferase UGT-18 protects worms from infection by pathogenic bacteria.

Altogether, this study identified mechanism by which olfactory neurons, activated by bacterial odours, control detoxification pathways resulting in enhanced survival of the host (Figure 8F).

## DISCUSSION

In this study, we found that innate immunity pathways in *C. elegans* could be primed by olfactory neurons sensing microbial odours. We show that two odours, known to cause aversion responses in *C*. *elegans*, upregulate hundreds of genes comprising various components of innate immunity in worms. These pathways provide broader protection against subsequent infection from Gram-negative and Gram-positive bacteria and pathogenic yeast. A UDP-glucuronosyltransferase, encoded by *ugt-18*, is a major effector upregulated in the intestine via the action of AWB olfactory neurons. UGT-18 protects worms from five diverse pathogens, at least one of which is known to produce the toxin phenazine. These findings suggest previously unappreciated and broad roles of odorants in activating immune responses in animals.

Animals possess several pathways to undertake the detoxification of a wide range of toxins produced by various pathogens while protecting their own tissues from damage. *P*. *aeruginosa* infects and wide range of plants and vertebrate and invertebrate hosts. It produces several toxins, including exotoxin A, pyocyanin, pyochelin, phenazines, and elastases^31,33–35^. In response, human epithelial cells induce pro-inflammatory cytokines, oxidative stress pathways, and apoptosis, while *C*. *elegans* activates the PMK-1/p38 MAPK pathway and upregulates detoxification enzymes like superoxide dismutase (SODs), UGTs, and cytochrome P450 (CYPs) to counteract oxidative stress caused by phenazines and pyocyanin of *P*. *aeruginosa*^35–37^. Other pathigens produce their arsenal of cytotoxic molecules. *E. faecalis* produces cytolysin, gelatinase, and pore-forming toxins^38^, which in humans elicit complement activation and tissue inhibitors of metalloproteinases (TIMPs), while *C. elegans* employs its PMK-1 pathway, antimicrobial peptides (AMPs), and CYPs as a defence^37,39–41^. *S. enterica*, with its SPI-1 effectors and typhoid toxin, induces inflammatory responses in humans through NF-κB and cGAS-STING pathways^42–44^, whereas in *C. elegans*, it triggers intestinal stress responses mediated by DAF-16/FOXO signalling to induce CYPs and SODs, along with AMPs^10,45–49^. For *S. aureus*, alpha-toxin causes pore formation and haemolysis in humans, leading to cytokine release and neutrophil recruitment^50–52^, while *C. elegans* mounts a PMK-1-dependent AMP response to intestinal damage^37,53,54^. Lastly, *C. neoformans* utilises its polysaccharide capsule and melanin production to evade immunity. In response, human macrophages attempt phagocytosis aided by interferon-gamma (IFN-γ) production^55,56^, while in *C. elegans*, capsule-mediated evasion is counteracted through the upregulation of lysozymes^57^. Some of these signalling pathways, such as PMK-1/MAPK^58^ and DAF-16/FOXO^59,60^, and immune effectors, such as AMPs, are known to be neuronally regulated in *C*. *elegans*^59,61–63^. These and other toxins produced by pathogens are substrates for UDP-glucuronosyltransferase enzymes of *C*. *elegans*, and UGT-18 may be one such enzyme to detoxify known and unknown toxins of microbial origin. Overall, this study is the first of its kind to demonstrate neuronal regulation of detoxification enzymes, such as UGTs, SODs, and CYPs, through olfaction.

Does the nervous system of other animals control detoxification pathways? In *D*. *melanogaster*, the central nervous system regulates detoxification enzymes like CYPs through neuropeptides and neurotransmitters^64^, while in molluscs like *Lymnaea stagnalis*, neurons coordinate antioxidant responses during metal toxicity^65^. In vertebrates, the hypothalamic-pituitary-adrenal (HPA) axis can induce cytochrome P450 enzymes through glucocorticoids during stress^66,67^, and in zebrafish, cortisol release from the brain activates detoxification pathways in the liver^68^. In *C. elegans*, sensory neurons rely on neuroendocrine signalling, particularly via insulin-like peptides that activate the DAF-2 receptor and modulate DAF-16/FOXO to regulate detoxification genes, including CYPs, UGTs, and glutathione-S-transferases (GSTs)^59,69,70^. Additionally, neuronal signals control SKN-1/Nrf2, a transcription factor that activates phase II detoxification enzymes in response to xenobiotic stress^71^. Neuronal regulation offers several advantages, including rapid responses to toxins, systemic coordination across tissues, energy-efficient activation of detoxification mechanisms, integration of behavioural changes like toxin avoidance, and adaptability to chronic toxin exposure, ensuring enhanced survival in dynamic environments.

The involvement of the olfactory system in regulating detoxification pathways, uncovered in this study, is the first of its kind. Many of the genes we identified to be regulated by AWB activation are reported to be expressed in the intestine of worms, the site of infection by most pathogens (WormExp analysis; Supplementary Table S2). A possible explanation for how AWB neurons in the head of the worm regulate gene expression in the non-innervated intestine could be via neuropeptide signalling. Neuropeptide transmission in *C*. *elegans* involves a complex network of signalling pathways that regulate various physiological processes, including behaviour, development, and stress responses^72^. The predominantly expressed neuropeptides of the AWB neurons include NLP-3 and NLP-9^73–75^. Neither of these neuropeptides has a known immune-related function. Similarly, NPR-42, the cognate receptor for NLP-3^76^, is also not well characterised with regard to immunity. Therefore, dissecting their roles in AWB-mediated immune response would open up exciting avenues of research, particularly in helping us understand the larger roles of neuropeptide signalling in organismal health and immunity.

This study highlights a hitherto unknown role of olfaction in host immunity. Olfactory GPCRs in AWB neurons likely function as non-canonical pathogen recognition receptors involved in “smelling” pathogens. Their identification will aim in understanding the role of olfaction in host-pathogen interaction, not only from the perspective of the worm but also from that of higher-order animals. For example, the human genome encodes nearly 800 GPCRs, of which roughly half are olfactory GPCRs (GPCRs_olf_)^77,78^. Many of these GPCRs_olf_ are expressed in non-nasal epithelial tissues, such as the lung, heart, kidney, and gut^79,80^. Recent studies have implicated non-nasal epithelial GPCRs_olf_ in sugar metabolism, sperm chemotaxis, and cell adhesion and migration^81–83^. Notably, several recent reports highlight the significance of the gut microbiota in mediating the gut-brain axis; this axis regulates issues related to human health, including the age-dependent decline in immunity, irritable bowel syndrome, cognition, anxiety, depression, reward/addiction pathway, metabolism, and eating disorders^84–89^. It is plausible that gut-microbiota-produced volatiles d mediate the gut-microbiota-brain axis through the GPCRs_olf_ in the gut epithelium. Studying the roles of olfactory GPCRs in the gut-microbiota-brain axis will advance research on metabolic and mental health disorders.

## ACKNOWLEDGEMENTS

Some *C. elegans* strains were provided by the CGC, which is funded by the NIH Office of Infrastructure Programs (P40 OD01440). SRV was supported by SRF fellowship from the Council for Scientific and Industrial Research (CSIR), India. We would like to thank Dr. Kavita Babu, Indian Institute of Science, for the discussions. This work was partly supported by the Wellcome Trust/DBT India Alliance Senior Fellowship (Grant no. IA/S/21/1/505655) and Royal Society Wolfson Fellowship (RSWF\R1\231005) to VS.

## AUTHOR CONTRIBUTIONS

Conceptualisation, SRV and VS; Methodology, SRV and VS; Formal Analysis, SRV and VS; Investigation, SRV and VS; Resources, VS; Writing Original Draft, SRV and VS; Writing, Review & Editing, SRV and VS; Fund Acquisition, VS; Supervision, VS.

## DATA AVAILABILITY

Fastq files for RNA-seq are available at NCBI (GEO Project ID GSE338878).

## RESOURCE AVAILABILITY

Further information and requests for resources or reagents should be directed to the lead contact. Reagents generated in this study are mentioned in Table S2.

## SUPPLEMENTAL INFORMATION

Table S1. Survival curve statistics.

Table S2. Transcriptome analysis and WormExp results.

Table S3. List of *C*. *elegans* strains used in this study.

## Materials and Methods

### *C*. *elegans* strains and maintenance

All *C*. *elegans* strains used in this study were maintained at 20°C as hermaphrodites on the nematode growth media (NGM; 0.3% NaCl, 0.25% peptone, 1 mM KPO_4_ buffer, 1 mM MgSO_4_, 1 mM CaCl_2_, 1.7% agar, 5 μg/mL cholesterol in absolute ethanol, and 50 μg/mL streptomycin in 1L double distilled water). The NGM plates were seeded with overnight cultures of *Escherichia coli* OP50. The wild-type N2 (Bristol) strain was used as the control. All strains used in this study are listed in Table S3.

### Bacterial strains and growth

*P*. *aeruginosa* PA14 was grown in LB broth overnight at 37°C, and 50 µl of the overnight culture was spread on slow killing (SK) agar plates^35^. The plates were then incubated at 37°C for 12 hours for *C. elegans* survival assays. *E*. *faecalis* OG1RF was grown in Brain Heart Infusion (BHI) broth supplemented with 50 µg/ml of gentamycin at 37°C for 8 hours and 50 µl of the culture was spread on BHI agar plates supplemented with 50 µg/ml of gentamycin. The plates were then incubated at 37°C for 12 hours for *C. elegans* survival assays^39^. *C*. *neoformans* H99α was grown in Yeast-extract Peptone Dextrose (YPD) broth for 12 hours at 25°C and 50 µl of the culture was spread on BHI agar plates. The plates were then incubated at 25°C for an additional 12 hours for *C. elegans* survival assays^57^. *S*. *enterica* was grown in LB broth overnight at 37°C and 50 µl of the overnight culture was spread on NGM plates. The plates were then incubated at 37°C for 12 hours for *C. elegans* survival assays^46^. *S*. *aureus* was grown in BHI broth for 5-6 hours at 37°C and 50 µl of the overnight culture was spread on BHI plates. The plates were then incubated at 37°C for 12 hours for *C. elegans* survival assays^53^. *E. coli* OP50 was grown in LB broth supplemented with 50 µg/ml of streptomycin at 37°C overnight. 200 µl of the culture was spotted on NGM plates for the maintenance of *C. elegans*^92^.

### Construction of transgenic strains

VSL2301 strain ((syIs371 [15xUAS::?*pes-10*::*HisCl1*::SL2::GFP::*let-858* 3’UTR + *unc- 122p*::GFP) + (syIs666 [*str-1p*::NLS::GAL4(sk)::VP64::*let-858* 3’UTR + *unc-122p*::RFP])) was created by crossing PS7199 [15xUAS::Δpes-10::HisCl1::SL2::GFP::let-858 3’UTR + *unc- 122p*::GFP] males with PS8565 [*str-1p*::NLS::GAL4(sk)::VP64::let-858 3’UTR + *unc- 122p*::RFP] hermaphrodites as per the standard protocol^107^. This strain was used as histamine responsive AWB neuron.

To create the *ugt-*18 reporter strain VSL2401 svEx[*ugt-18p*::GFP + *myo-2p*::mCherry], a 2009 bp fragment containing the *ugt-18* promoter was amplified from the genomic DNA of WT worms using the primers 5’- GATGCATGCGTTCCGAGTGTAGTCAACAA-3’ and 5’- GTACTGCAGGATCGGAGTAATCTTAACTTG-3’. This fragment was cloned into the pPD95_75 plasmid to yield pPD95_75_*ugt-18p*::GFP. Then, 50 ng/µL of pPD95_75_ *ugt- 18p*::GFP and 5 ng/µL of pCFJ90 (*myo-2p*::mCherry::*unc54*-UTR) were microinjected into WT worms to create VSL2401.

### Survival assays on pathogens

The survival assays with the pathogens were performed as previously described^17,35,46,53,57,93^. 50-60 age-synchronised young adults were transferred to the pathogen plates and the plates were incubated at 25°C till the entire population of worms was dead. The plates were scored for live and dead worms twice a day. Kaplan-Meier plots for survival were plotted and analysed by Log-Rank test. At least 3 biological repeats were performed, and the statistics for each biological repeat for survival assays are presented in Table S1.

### Ligand-mediated neuronal activation

For ligand-mediated activation of the AWB neurons in WT worms, either 1-undecene or 2- nonanone was used^19,22^. Diacetyl, sensed by the AWA neurons in WT, was used as a control. Furthermore, in the CX3877 strain, expressing the diacetyl receptor (ODR-10) in the AWB neuron, diacetyl was used to activate AWB. Briefly, age-synchronised worms were grown on NGM-OP50 plates. When the worms reached the L4 stage, they were inverted onto an SK plate containing the spots of the test volatile, sealed using parafilm, and incubated at 25°C for 90 minutes. For RNASeq experiments, worms were exposed to 1 μL of 2-nonanone (Sigma Aldrich, Burlington, MA, USA) or 4 spots of 1-undecene (3 μL each; Sigma Aldrich). For pre-exposure experiments to prime the immune system, worms were exposed to 0.5 μL of 2- nonanone, 3 μL of 1-undecene, or 3 μL of diacetyl (Sigma Aldrich). For all experiments, worms were allowed to recover at 20°C for 2 hours after the removal of the volatile cue. After the recovery period, RNA isolation or survival assays were performed.

### Histamine-mediated neuronal silencing

In this study, the VSL2301 strain ((syIs371 [15xUAS::?*pes-10*::*HisCl1*::SL2::GFP::*let-858* 3’UTR + *unc-122p*::GFP) + (syIs666 [*str-1p*::NLS::GAL4(sk)::VP64::*let-858* 3’UTR + *unc-122p*::RFP])), expressing the *Drosophila* HisCl1 receptor in the AWB neurons, was used for silencing the AWB neurons. Previously reported protocols with slight modifications were used for histamine mediated silencing^23^. Briefly, worms were placed on NGM-OP50 plates supplemented with 10 mM histamine dichloride (Sigma Aldrich) for 4 or 8 hours prior to the assay. Silencing of the AWB neuron was confirmed by evaluating chemotaxis. Prolonged silencing of the AWB neurons during *P. aeruginosa* infection were performed on NGM-PA14 plates containing 15 mM histamine dichloride.

### Chemotaxis assay

All chemotaxis assays were performed on 90 mm plates of buffered agar (1 mM CaCl_2_, 1 mM MgSO_4_, 25 mM KPO_4_, and 2% agar). Freshly poured buffered agar plates were dried for 50- 60 minutes with open lids inside a laminar airflow and used for chemotaxis after 24-36 hours. The test chemical (either absolute chemical or diluted in an appropriate solvent) was spotted at one end of the 90 mm plate, while the solvent control was spotted at the diagonally opposite end. 2 μL sodium azide was spotted next to test and control spots to paralyse worms that have moved towards either end. Approximately 50-60 young adult worms were washed thrice in S-basal buffer (5.85 g NaCl, 1 g K_2_HPO_4_, 6 g KH_2_PO_4_, 1 mL cholesterol (5 mg/mL in ethanol), double distilled water to 1 L) and placed at the centre of the assay plate. The plates were sealed with parafilm and incubated at 25°C for 2 hours. Worms on either side of the plates were counted, and the chemotaxis index (CI) was calculated as follows^19,94^:

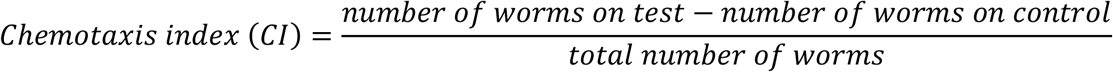

### RNA isolation

Synchronised WT worms were grown on NGM-OP50 and exposed to odours as described. Roughly 1,000 L4 worms were harvested and washed three times in M9 buffer (3 g of KH_2_PO_4_, 6 g of Na_2_HPO_4_, 5 g NaCl, and 1 mL of 1 M MgSO_4_ in 1 L double distilled water). 800 µL of Qiazol reagent (Qiagen, Germany) was added to the wprm pellet and and frozen at -80°C. The frozen worms were thawed at 32°C in a water bath, vortexed for 90 seconds, and flash-frozen in liquid nitrogen for 90 seconds. This was repeated six times. The lysate was then used for RNA extraction using the Qiagen Universal RNA isolation kit (Qiagen, Germany) as per the manufacturer’s protocol. Three biological replicates were prepared for each sample.

### mRNA library preparation, RNA sequencing, and analysis

The ready-to-sequence library was prepared using the NEBNext Ultra II Directional RNA Library Prep Kit for Illumina (NEB #E7765). The libraries of the samples were then subjected to 150-base pairs paired-end Illumina sequencing, resulting in 35-45 million reads per sample at Clevergene Biocorp. Pvt. Ltd. The raw data obtained from the sequencing were processed using the RNASeq analysis pipeline described previously^95^. HISAT2 was used to map the RNASeq reads with the latest *C. elegans* reference genome (WS280). The output was in the form of a Sequence Alignment Map (SAM). The SAM format was converted to its binary form, BAM, using SAMTOOLS and was then used as the input for assembling the transcripts using StringTie to quantify the levels of expressed genes and obtain Fragments Per Kilobase of exon per Million maps read (FPKM) values. The assembly was merged into a singular Gene Transfer Format (GTF) to facilitate comparison with the reference annotation in the same format.

The fold change was calculated using the formula:

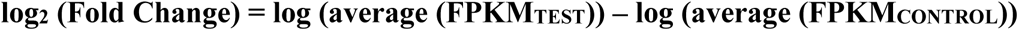

Genes that were upregulated and downregulated above a cut-off fold change of 2-fold with a *P*-value of less than 0.05 were shortlisted. Volcano plots were prepared using RStudio. The unique WormBase IDs of the shortlisted genes were used as inputs, and the genes reported to be enriched for gene ontology (GO) of biological processes were obtained from DAVID and WormBase^96–99^. An additional gene enrichment analysis was performed using the Worm Exp tool (https://wormexp.zoologie.uni-kiel.de/wormexp/)^100^. We performed GO studies on the upregulated genes using the DAVID and WormBase tools^97–99,101–104^.

### Quantitative real-time-PCR

Quantitative real-time PCR (qRT-PCR) was performed to evaluate the differentially regulated genes under various treatments. Total RNA was isolated as described above. Genomic DNA contamination was removed using DNase I (Thermo Fischer, Waltham, MA, USA) in accordance with the manufacturer’s protocol. Next, cDNA was prepared from the DNA-depleted total RNA using the iScript cDNA synthesis kit (Bio-Rad, Hercules, CA, USA). qRT-PCR was performed on the cDNA using iTaq Universal SYBR green supermix (Bio-Rad) on an Applied Biosystem Quant 3 Studio set-up^105^. Briefly, 50 ng of the cDNA was mixed with iTaq Universal SYBR green supermix (2x), as well as the forward and reverse primers, to prepare the master mix. 10 µL of this mix was aliquoted into clear-bottom 96-well plates, and qRT-PCR was performed as per the manufacturer’s protocol. Then, the obtained C_T_ values were used to calculate the fold change based on the Livak 2^−ΔΔ*CT*^ method^106^. A lower C_T_ value indicates higher target mRNA expression, while a higher value indicates lower expression. All target C_T_ values were normalised to that of *act-1* (Actin). At least three biological replicates were performed. Primer sequences for qRT-PCR are available upon request.

### Gene knockdown using RNAi

Systemic RNAi by feeding was performed as previously described^91,108^ using the *E*. *coli* HT115(DE3) RNAi clones available in the Ahringer library ^90^. Bacteria carrying RNAI clone against the gene of interest was grown at 37°C in LB broth supplemented with 50 µg/mL carbenicillin. 200 µL of the culture was then spotted on NGM plates containing 50 µg/mL carbenicillin and 5 mM Isopropyl β-D-1-thiogalactopyranoside (IPTG, SRL Chemicals, India) and incubated at 25°C overnight. Worms were allowed to develop on these plates at 20°C until further use. As a control, HT115(DE3) bacteria containing an empty L4440 plasmid was used.

### Phenazine toxicity assay

The phenazine toxicity assay was performed as previously described^30^. 30 young adult worms were transferred to NGM-OP50 plates supplemented with 180 µM phenazine (Sigma Aldrich) solubilized in DMSO. NGM-OP50 plates with DMSOwere used as the control. The number of live worms at every time point was recorded at each time point. For subsequent assays live worms were counted at 9 hours after phenazine exposure for various genotypes, RNAi and odour exposure conditions in at least three biological replicate.

### Fluorescence microscopy

Control or test worms were synchronised on NGM-OP50 until adulthood. Adult worms were then subjected to control or odour treatments, washed thrice in M9 buffer, mounted on a glass slide with 2% agarose padding, and anaesthetised using 0.5 µL of 1 M sodium azide. The mounted worms were then visualised and imaged under a fluorescence microscope (Olympus IX81, Tokyo, Japan). Comparisons were done using the ImageJ software.

### Statistical analyses

Statistical analyses for all assays were performed using GraphPad PRISM 8.0. Kaplan-Meier survival curves for survival assays on pathogens were plotted, and statistical analyses relied on the Log-Rank test. Analysis of variance (ANOVA) followed by a post-hoc Dunnet test with adjusted *P*-values was used for multiple comparisons, while Student’s unpaired t-test with Welch’s correction was used for one-on-one comparisons. Statistical significance was set at *P* < 0.05. \**P* < 0.05; \*\**P* < 0.01; \*\*\**P* < 0.001; \*\*\*\**P* < 0.0001.

## REFERENCES

1. Akira, S., Uematsu, S., and Takeuchi, O. (2006). Pathogen recognition and innate immunity. Cell 124, 783–801. 10.1016/j.cell.2006.02.015.

2. Kumagai, Y., and Akira, S. (2010). Identification and functions of pattern-recognition receptors. J Allergy Clin Immunol 125, 985–992. 10.1016/j.jaci.2010.01.058.

3. Janeway, C.A., Jr., and Medzhitov, R. (2002). Innate immune recognition. Annual review of immunology 20, 197–216. 10.1146/annurev.immunol.20.083001.084359.

4. Medzhitov, R., and Janeway, C., Jr. (2000). Innate immune recognition: mechanisms and pathways. Immunol Rev 173, 89–97. 10.1034/j.1600-065x.2000.917309.x.

5. Pukkila-Worley, R., and Ausubel, F.M. (2012). Immune defense mechanisms in the Caenorhabditis elegans intestinal epithelium. Curr Opin Immunol 24, 3–9. 10.1016/j.coi.2011.10.004.

6. Lemaitre, B., and Hoffmann, J. (2007). The host defense of Drosophila melanogaster. Annual review of immunology 25, 697–743. 10.1146/annurev.immunol.25.022106.141615.

7. Hansen, J.D., Vojtech, L.N., and Laing, K.J. (2011). Sensing disease and danger: a survey of vertebrate PRRs and their origins. Dev Comp Immunol 35, 886–897. 10.1016/j.dci.2011.01.008.

8. Lemaitre, B., Nicolas, E., Michaut, L., Reichhart, J.M., and Hoffmann, J.A. (1996). The dorsoventral regulatory gene cassette spatzle/Toll/cactus controls the potent antifungal response in Drosophila adults. Cell 86, 973-983. 10.1016/s0092-8674(00)80172-5.

9. Engelmann, I., and Pujol, N. (2010). Innate immunity in C. elegans. Advances in experimental medicine and biology 708, 105–121. 10.1007/978-1-4419-8059-5_6.

10. Tenor, J.L., and Aballay, A. (2008). A conserved Toll-like receptor is required for Caenorhabditis elegans innate immunity. EMBO Rep 9, 103-109. 10.1038/sj.embor.7401104.

11. Pradel, E., Zhang, Y., Pujol, N., Matsuyama, T., Bargmann, C.I., and Ewbank, J.J. (2007). Detection and avoidance of a natural product from the pathogenic bacterium Serratia marcescens by Caenorhabditis elegans. Proceedings of the National Academy of Sciences of the United States of America 104, 2295–2300. 10.1073/pnas.0610281104.

12. Pujol, N., Link, E.M., Liu, L.X., Kurz, C.L., Alloing, G., Tan, M.W., Ray, K.P., Solari, R., Johnson, C.D., and Ewbank, J.J. (2001). A reverse genetic analysis of components of the Toll signaling pathway in Caenorhabditis elegans. Curr Biol 11, 809–821. 10.1016/s0960-9822(01)00241-x.

13. Sowa, J.N., Jiang, H., Somasundaram, L., Tecle, E., Xu, G., Wang, D., and Troemel, E.R. (2020). The Caenorhabditis elegans RIG-I Homolog DRH-1 Mediates the Intracellular Pathogen Response upon Viral Infection. Journal of virology 94. 10.1128/JVI.01173-19.

14. Miltsch, S.M., Seeberger, P.H., and Lepenies, B. (2014). The C-type lectin-like domain containing proteins Clec-39 and Clec-49 are crucial for Caenorhabditis elegans immunity against Serratia marcescens infection. Dev Comp Immunol 45, 67–73. 10.1016/j.dci.2014.02.002.

15. Venkatesh, S.R., and Singh, V. (2021). G protein-coupled receptors: The choreographers of innate immunity in Caenorhabditis elegans. PLoS Pathog 17, e1009151. 10.1371/journal.ppat.1009151.

16. Miller, E.V., Grandi, L.N., Giannini, J.A., Robinson, J.D., and Powell, J.R. (2015). The Conserved G-Protein Coupled Receptor FSHR-1 Regulates Protective Host Responses to Infection and Oxidative Stress. PLoS One 10, e0137403. 10.1371/journal.pone.0137403.

17. Venkatesh, S.R., Gupta, A., and Singh, V. (2023). Amphid sensory neurons of Caenorhabditis elegans orchestrate its survival from infection with broad classes of pathogens. Life Sci Alliance 6. 10.26508/lsa.202301949.

18. Meisel, J.D., Panda, O., Mahanti, P., Schroeder, F.C., and Kim, D.H. (2014). Chemosensation of bacterial secondary metabolites modulates neuroendocrine signaling and behavior of C. elegans. Cell 159, 267-280. 10.1016/j.cell.2014.09.011.

19. Prakash, D., Ms, A., Radhika, B., Venkatesan, R., Chalasani, S.H., and Singh, V. (2021). 1-Undecene from Pseudomonas aeruginosa is an olfactory signal for flight-or-fight response in Caenorhabditis elegans. The EMBO journal 40, e106938. 10.15252/embj.2020106938.

20. Sengupta, P. (2013). The belly rules the nose: feeding state-dependent modulation of peripheral chemosensory responses. Curr Opin Neurobiol 23, 68–75. 10.1016/j.conb.2012.08.001.

21. Srinivasan, J., Kaplan, F., Ajredini, R., Zachariah, C., Alborn, H.T., Teal, P.E., Malik, R.U., Edison, A.S., Sternberg, P.W., and Schroeder, F.C. (2008). A blend of small molecules regulates both mating and development in Caenorhabditis elegans. Nature 454, 1115–1118. 10.1038/nature07168.

22. Troemel, E.R., Kimmel, B.E., and Bargmann, C.I. (1997). Reprogramming chemotaxis responses: sensory neurons define olfactory preferences in C. elegans. Cell 91, 161–169. 10.1016/s0092-8674(00)80399-2.

23. Pokala, N., Liu, Q., Gordus, A., and Bargmann, C.I. (2014). Inducible and titratable silencing of Caenorhabditis elegans neurons in vivo with histamine-gated chloride channels. Proceedings of the National Academy of Sciences of the United States of America 111, 2770–2775. 10.1073/pnas.1400615111.

24. Tang, D., Chen, M., Huang, X., Zhang, G., Zeng, L., Zhang, G., Wu, S., and Wang, Y. (2023). SRplot: A free online platform for data visualization and graphing. PLoS One 18, e0294236. 10.1371/journal.pone.0294236.

25. Dasgupta, M., Shashikanth, M., Gupta, A., Sandhu, A., De, A., Javed, S., and Singh, V. (2020). NHR-49 Transcription Factor Regulates Immunometabolic Response and Survival of Caenorhabditis elegans during Enterococcus faecalis Infection. Infect Immun 88. 10.1128/IAI.00130-20.

26. Tukey, R.H., and Strassburg, C.P. (2000). Human UDP-glucuronosyltransferases: metabolism, expression, and disease. Annu Rev Pharmacol Toxicol 40, 581–616. 10.1146/annurev.pharmtox.40.1.581.

27. Perreault, M., Bialek, A., Trottier, J., Verreault, M., Caron, P., Milkiewicz, P., and Barbier, O. (2013). Role of glucuronidation for hepatic detoxification and urinary elimination of toxic bile acids during biliary obstruction. PLoS One 8, e80994. 10.1371/journal.pone.0080994.

28. Fontaine, P., and Choe, K. (2018). The transcription factor SKN-1 and detoxification gene ugt-22 alter albendazole efficacy in Caenorhabditis elegans. Int J Parasitol Drugs Drug Resist 8, 312–319. 10.1016/j.ijpddr.2018.04.006.

29. Sharma, N., Au, V., Martin, K., Edgley, M.L., Moerman, D., Mains, P.E., and Gilleard, J.S. (2024). Multiple UDP glycosyltransferases modulate benzimidazole drug sensitivity in the nematode Caenorhabditis elegans in an additive manner. Int J Parasitol. 10.1016/j.ijpara.2024.05.003.

30. Asif, M.Z., Nocilla, K.A., Ngo, L., Shah, M., Smadi, Y., Hafeez, Z., Parnes, M., Manson, A., Glushka, J.N., Leach, F.E., 3rd, and Edison, A.S. (2024). Role of UDP-Glycosyltransferase (ugt) Genes in Detoxification and Glycosylation of 1- Hydroxyphenazine (1-HP) in Caenorhabditis elegans. Chem Res Toxicol *37*, 590-599. 10.1021/acs.chemrestox.3c00410.

31. Cezairliyan, B., Vinayavekhin, N., Grenfell-Lee, D., Yuen, G.J., Saghatelian, A., and Ausubel, F.M. (2013). Identification of Pseudomonas aeruginosa phenazines that kill Caenorhabditis elegans. PLoS Pathog *9*, e1003101. 10.1371/journal.ppat.1003101.

32. Wilson, R., Sykes, D.A., Watson, D., Rutman, A., Taylor, G.W., and Cole, P.J. (1988). Measurement of Pseudomonas aeruginosa phenazine pigments in sputum and assessment of their contribution to sputum sol toxicity for respiratory epithelium. Infect Immun 56, 2515-2517. 10.1128/iai.56.9.2515-2517.1988.

33. Qin, S., Xiao, W., Zhou, C., Pu, Q., Deng, X., Lan, L., Liang, H., Song, X., and Wu, M. (2022). Pseudomonas aeruginosa: pathogenesis, virulence factors, antibiotic resistance, interaction with host, technology advances and emerging therapeutics. Signal Transduct Target Ther 7, 199. 10.1038/s41392-022-01056-1.

34. Wood, S.J., Goldufsky, J.W., Seu, M.Y., Dorafshar, A.H., and Shafikhani, S.H. (2023). Pseudomonas aeruginosa Cytotoxins: Mechanisms of Cytotoxicity and Impact on Inflammatory Responses. Cells 12. 10.3390/cells12010195.

35. Tan, M.W., Mahajan-Miklos, S., and Ausubel, F.M. (1999). Killing of Caenorhabditis elegans by Pseudomonas aeruginosa used to model mammalian bacterial pathogenesis. Proceedings of the National Academy of Sciences of the United States of America Sc, 715-720. 10.1073/pnas.96.2.715.

36. Kim, D.H., Feinbaum, R., Alloing, G., Emerson, F.E., Garsin, D.A., Inoue, H., Tanaka-Hino, M., Hisamoto, N., Matsumoto, K., Tan, M.W., and Ausubel, F.M. (2002). A conserved p38 MAP kinase pathway in Caenorhabditis elegans innate immunity. Science 2S*7*, 623-626. 10.1126/science.1073759.

37. Irazoqui, J.E., Troemel, E.R., Feinbaum, R.L., Luhachack, L.G., Cezairliyan, B.O., and Ausubel, F.M. (2010). Distinct pathogenesis and host responses during infection of C. elegans by P. aeruginosa and S. aureus. PLoS Pathog *6*, e1000982. 10.1371/journal.ppat.1000982.

38. Fiore, E., Van Tyne, D., and Gilmore, M.S. (2019). Pathogenicity of Enterococci. Microbiol Spectr 7. 10.1128/microbiolspec.GPP3-0053-2018.

39. Garsin, D.A., Sifri, C.D., Mylonakis, E., Qin, X., Singh, K.V., Murray, B.E., Calderwood, S.B., and Ausubel, F.M. (2001). A simple model host for identifying Gram-positive virulence factors. Proceedings of the National Academy of Sciences of the United States of America 98, 10892–10897. 10.1073/pnas.191378698.

40. Sifri, C.D., Mylonakis, E., Singh, K.V., Qin, X., Garsin, D.A., Murray, B.E., Ausubel, F.M., and Calderwood, S.B. (2002). Virulence effect of Enterococcus faecalis protease genes and the quorum-sensing locus fsr in Caenorhabditis elegans and mice. Infect Immun 70, 5647–5650. 10.1128/IAI.70.10.5647-5650.2002.

41. Yuen, G.J., and Ausubel, F.M. (2018). Both live and dead Enterococci activate Caenorhabditis elegans host defense via immune and stress pathways. Virulence 9, 683-699. 10.1080/21505594.2018.1438025.

42. Fabrega, A., and Vila, J. (2013). Salmonella enterica serovar Typhimurium skills to succeed in the host: virulence and regulation. Clin Microbiol Rev 26, 308-341. 10.1128/CMR.00066-12.

43. Anderson, C.J., and Kendall, M.M. (2017). Salmonella enterica Serovar Typhimurium Strategies for Host Adaptation. Front Microbiol 8, 1983. 10.3389/fmicb.2017.01983.

44. Haraga, A., Ohlson, M.B., and Miller, S.I. (2008). Salmonellae interplay with host cells. Nat Rev Microbiol 6, 53-66. 10.1038/nrmicro1788.

45. Aballay, A., Drenkard, E., Hilbun, L.R., and Ausubel, F.M. (2003). Caenorhabditis elegans innate immune response triggered by Salmonella enterica requires intact LPS and is mediated by a MAPK signaling pathway. Curr Biol 13, 47–52. 10.1016/s0960-9822(02)01396-9.

46. Aballay, A., Yorgey, P., and Ausubel, F.M. (2000). Salmonella typhimurium proliferates and establishes a persistent infection in the intestine of Caenorhabditis elegans. Curr Biol 10, 1539–1542. 10.1016/s0960-9822(00)00830-7.

47. Aballay, A., and Ausubel, F.M. (2001). Programmed cell death mediated by ced-3 and ced-4 protects Caenorhabditis elegans from Salmonella typhimurium-mediated killing. Proceedings of the National Academy of Sciences of the United States of America 98, 2735–2739. 10.1073/pnas.041613098.

48. Alegado, R.A., and Tan, M.W. (2008). Resistance to antimicrobial peptides contributes to persistence of Salmonella typhimurium in the C. elegans intestine. Cell Microbiol 10, 1259–1273. 10.1111/j.1462-5822.2008.01124.x.

49. Sem, X., and Rhen, M. (2012). Pathogenicity of Salmonella enterica in Caenorhabditis elegans relies on disseminated oxidative stress in the infected host. PLoS One 7, e45417. 10.1371/journal.pone.0045417.

50. Bhakdi, S., and Tranum-Jensen, J. (1991). Alpha-toxin of Staphylococcus aureus. Microbiol Rev 55, 733–751. 10.1128/mr.55.4.733-751.1991.

51. Cheung, G.Y.C., Bae, J.S., and Otto, M. (2021). Pathogenicity and virulence of Staphylococcus aureus. Virulence 12, 547–569. 10.1080/21505594.2021.1878688.

52. Otto, M. (2014). Staphylococcus aureus toxins. Curr Opin Microbiol 17, 32–37. 10.1016/j.mib.2013.11.004.

53. Sifri, C.D., Begun, J., Ausubel, F.M., and Calderwood, S.B. (2003). Caenorhabditis elegans as a model host for Staphylococcus aureus pathogenesis. Infect Immun 71, 2208–2217. 10.1128/IAI.71.4.2208-2217.2003.

54. Wani, K.A., Goswamy, D., Taubert, S., Ratnappan, R., Ghazi, A., and Irazoqui, J.E. (2021). NHR-49/PPAR-alpha and HLH-30/TFEB cooperate for C. elegans host defense via a flavin-containing monooxygenase. eLife 10. 10.7554/eLife.62775.

55. Zaragoza, O., Rodrigues, M.L., De Jesus, M., Frases, S., Dadachova, E., and Casadevall, A. (2009). The capsule of the fungal pathogen Cryptococcus neoformans. Adv Appl Microbiol 68, 133–216. 10.1016/S0065-2164(09)01204-0.

56. Rodrigues, M.L., Alviano, C.S., and Travassos, L.R. (1999). Pathogenicity of Cryptococcus neoformans: virulence factors and immunological mechanisms. Microbes Infect 1, 293–301. 10.1016/s1286-4579(99)80025-2.

57. Mylonakis, E., Ausubel, F.M., Perfect, J.R., Heitman, J., and Calderwood, S.B. (2002). Killing of Caenorhabditis elegans by Cryptococcus neoformans as a model of yeast pathogenesis. Proceedings of the National Academy of Sciences of the United States of America SS, 15675-15680. 10.1073/pnas.232568599.

58. Foster, K.J., Cheesman, H.K., Liu, P., Peterson, N.D., Anderson, S.M., and Pukkila-Worley, R. (2020). Innate Immunity in the C. elegans Intestine Is Programmed by a Neuronal Regulator of AWC Olfactory Neuron Development. Cell Rep 31, 107478. 10.1016/j.celrep.2020.03.042.

59. Kawli, T., and Tan, M.W. (2008). Neuroendocrine signals modulate the innate immunity of Caenorhabditis elegans through insulin signaling. Nat Immunol 9, 1415-1424. 10.1038/ni.1672.

60. Evans, E.A., Kawli, T., and Tan, M.W. (2008). Pseudomonas aeruginosa suppresses host immunity by activating the DAF-2 insulin-like signaling pathway in Caenorhabditis elegans. PLoS Pathog 4, e1000175. 10.1371/journal.ppat.1000175.

61. Styer, K.L., Singh, V., Macosko, E., Steele, S.E., Bargmann, C.I., and Aballay, A. (2008). Innate immunity in Caenorhabditis elegans is regulated by neurons expressing NPR-1/GPCR. Science 322, 460–464. 10.1126/science.1163673.

62. Wibisono, P., and Sun, J. (2021). Neuro-immune communication in C. elegans defense against pathogen infection. Curr Res Immunol 2, 60–65. 10.1016/j.crimmu.2021.04.002.

63. Zugasti, O., and Ewbank, J.J. (2009). Neuroimmune regulation of antimicrobial peptide expression by a noncanonical TGF-beta signaling pathway in Caenorhabditis elegans epidermis. Nat Immunol 10, 249–256. 10.1038/ni.1700.

64. Baldwin, S.R., Mohapatra, P., Nagalla, M., Sindvani, R., Amaya, D., Dickson, H.A., and Menuz, K. (2021). Identification and characterization of CYPs induced in the Drosophila antenna by exposure to a plant odorant. Sci Rep 11, 20530. 10.1038/s41598-021-99910-9.

65. Gnatyshyna, L., Khoma, V., Martinyuk, V., Matskiv, T., Pedrini-Martha, V., Niederwanger, M., Stoliar, O., and Dallinger, R. (2023). Sublethal cadmium exposure in the freshwater snail Lymnaea stagnalis meets a deficient, poorly responsive metallothionein system while evoking oxidative and cellular stress. Comp Biochem Physiol C Toxicol Pharmacol 2c*3*, 109490. 10.1016/j.cbpc.2022.109490.

66. Stephens, M.A., and Wand, G. (2012). Stress and the HPA axis: role of glucocorticoids in alcohol dependence. Alcohol Res 34, 468–483.

67. Burford, N.G., Webster, N.A., and Cruz-Topete, D. (2017). Hypothalamic-Pituitary-Adrenal Axis Modulation of Glucocorticoids in the Cardiovascular System. Int J Mol Sci 18. 10.3390/ijms18102150.

68. Alsop, D., and Vijayan, M.M. (2009). Molecular programming of the corticosteroid stress axis during zebrafish development. Comp Biochem Physiol A Mol Integr Physiol 153, 49–54. 10.1016/j.cbpa.2008.12.008.

69. Murphy, C.T., McCarroll, S.A., Bargmann, C.I., Fraser, A., Kamath, R.S., Ahringer, J., Li, H., and Kenyon, C. (2003). Genes that act downstream of DAF-16 to influence the lifespan of Caenorhabditis elegans. Nature 424, 277–283. 10.1038/nature01789.

70. Troemel, E.R., Chu, S.W., Reinke, V., Lee, S.S., Ausubel, F.M., and Kim, D.H. (2006). p38 MAPK regulates expression of immune response genes and contributes to longevity in C. elegans. PLoS Genet 2, e183. 10.1371/journal.pgen.0020183.

71. An, J.H., and Blackwell, T.K. (2003). SKN-1 links C. elegans mesendodermal specification to a conserved oxidative stress response. Genes Dev 17, 1882–1893. 10.1101/gad.1107803.

72. Li, C., and Kim, K. (2008). Neuropeptides. WormBook, 1-36. 10.1895/wormbook.1.142.1.

73. Nathoo, A.N., Moeller, R.A., Westlund, B.A., and Hart, A.C. (2001). Identification of neuropeptide-like protein gene families in Caenorhabditiselegans and other species. Proceedings of the National Academy of Sciences of the United States of America 98, 14000–14005. 10.1073/pnas.241231298.

74. Hammarlund, M., Hobert, O., Miller, D.M., 3rd, and Sestan, N. (2018). The CeNGEN Project: The Complete Gene Expression Map of an Entire Nervous System. Neuron SS, 430-433. 10.1016/j.neuron.2018.07.042.

75. Taylor, S.R., Santpere, G., Weinreb, A., Barrett, A., Reilly, M.B., Xu, C., Varol, E., Oikonomou, P., Glenwinkel, L., McWhirter, R., et al. (2021). Molecular topography of an entire nervous system. Cell 184, 4329–4347 e4323. 10.1016/j.cell.2021.06.023.

76. Beets, I., Zels, S., Vandewyer, E., Demeulemeester, J., Caers, J., Baytemur, E., Courtney, A., Golinelli, L., Hasakiogullari, I., Schafer, W.R., et al. (2023). System-wide mapping of peptide-GPCR interactions in C. elegans. Cell Rep 42, 113058. 10.1016/j.celrep.2023.113058.

77. Mombaerts, P. (2004). Genes and ligands for odorant, vomeronasal and taste receptors. Nat Rev Neurosci 5, 263–278. 10.1038/nrn1365.

78. Trimmer, C., Keller, A., Murphy, N.R., Snyder, L.L., Willer, J.R., Nagai, M.H., Katsanis, N., Vosshall, L.B., Matsunami, H., and Mainland, J.D. (2019). Genetic variation across the human olfactory receptor repertoire alters odor perception. Proceedings of the National Academy of Sciences of the United States of America 116, 9475-9480. 10.1073/pnas.1804106115.

79. Flegel, C., Manteniotis, S., Osthold, S., Hatt, H., and Gisselmann, G. (2013). Expression profile of ectopic olfactory receptors determined by deep sequencing. PLoS One 8, e55368. 10.1371/journal.pone.0055368.

80. Shepard, B.D. (2021). The Sniffing Kidney: Roles for Renal Olfactory Receptors in Health and Disease. Kidney360 *2*, 1056–1062. 10.34067/KID.0000712021.

81. Griffin, C.A., Kafadar, K.A., and Pavlath, G.K. (2009). MOR23 promotes muscle regeneration and regulates cell adhesion and migration. Dev Cell 17, 649–661. 10.1016/j.devcel.2009.09.004.

82. Kim, K.S., Lee, I.S., Kim, K.H., Park, J., Kim, Y., Choi, J.H., Choi, J.S., and Jang, H.J. (2017). Activation of intestinal olfactory receptor stimulates glucagon-like peptide-1 secretion in enteroendocrine cells and attenuates hyperglycemia in type 2 diabetic mice. Sci Rep 7, 13978. 10.1038/s41598-017-14086-5.

83. Spehr, M., Gisselmann, G., Poplawski, A., Riffell, J.A., Wetzel, C.H., Zimmer, R.K., and Hatt, H. (2003). Identification of a testicular odorant receptor mediating human sperm chemotaxis. Science 2S*S*, 2054-2058. 10.1126/science.1080376.

84. Cryan, J.F., O’Riordan, K.J., Cowan, C.S.M., Sandhu, K.V., Bastiaanssen, T.F.S., Boehme, M., Codagnone, M.G., Cussotto, S., Fulling, C., Golubeva, A.V., et al. (2019). The Microbiota-Gut-Brain Axis. Physiol Rev SS, 1877-2013. 10.1152/physrev.00018.2018.

85. Long-Smith, C., O’Riordan, K.J., Clarke, G., Stanton, C., Dinan, T.G., and Cryan, J.F. (2020). Microbiota-Gut-Brain Axis: New Therapeutic Opportunities. Annu Rev Pharmacol Toxicol 60, 477–502. 10.1146/annurev-pharmtox-010919-023628.

86. de Theije, C.G., Wopereis, H., Ramadan, M., van Eijndthoven, T., Lambert, J., Knol, J., Garssen, J., Kraneveld, A.D., and Oozeer, R. (2014). Altered gut microbiota and activity in a murine model of autism spectrum disorders. Brain Behav Immun 37, 197–206. 10.1016/j.bbi.2013.12.005.

87. Farmer, A.D., and Aziz, Q. (2014). Mechanisms and management of functional abdominal pain. J R Soc Med 107, 347–354. 10.1177/0141076814540880.

88. Guigoz, Y., Dore, J., and Schiffrin, E.J. (2008). The inflammatory status of old age can be nurtured from the intestinal environment. Curr Opin Clin Nutr Metab Care 11, 13–20. 10.1097/MCO.0b013e3282f2bfdf.

89. Hou, K., Wu, Z.X., Chen, X.Y., Wang, J.Q., Zhang, D., Xiao, C., Zhu, D., Koya, J.B., Wei, L., Li, J., and Chen, Z.S. (2022). Microbiota in health and diseases. Signal Transduct Target Ther 7, 135. 10.1038/s41392-022-00974-4.

90. Kamath, R.S., and Ahringer, J. (2003). Genome-wide RNAi screening in Caenorhabditis elegans. Methods 30, 313–321. 10.1016/s1046-2023(03)00050-1.

91. Kamath, R.S., Martinez-Campos, M., Zipperlen, P., Fraser, A.G., and Ahringer, J. (2001). Effectiveness of specific RNA-mediated interference through ingested double-stranded RNA in Caenorhabditis elegans. Genome Biol 2, RESEARCH0002. 10.1186/gb-2000-2-1-research0002.

92. Brenner, S. (1974). The genetics of Caenorhabditis elegans. Genetics 77, 71–94.

93. Garsin, D.A., Villanueva, J.M., Begun, J., Kim, D.H., Sifri, C.D., Calderwood, S.B., Ruvkun, G., and Ausubel, F.M. (2003). Long-lived C. elegans daf-2 mutants are resistant to bacterial pathogens. Science 300, 1921. 10.1126/science.1080147.

94. Bargmann, C.I., Hartwieg, E., and Horvitz, H.R. (1993). Odorant-selective genes and neurons mediate olfaction in C. elegans. Cell 74, 515–527. 10.1016/0092-8674(93)80053-h.

95. Pertea, M., Kim, D., Pertea, G.M., Leek, J.T., and Salzberg, S.L. (2016). Transcript-level expression analysis of RNA-seq experiments with HISAT, StringTie and Ballgown. Nat Protoc 11, 1650–1667. 10.1038/nprot.2016.095.

96. Grove, C., Cain, S., Chen, W.J., Davis, P., Harris, T., Howe, K.L., Kishore, R., Lee, R., Paulini, M., Raciti, D., et al. (2018). Using WormBase: A Genome Biology Resource for Caenorhabditis elegans and Related Nematodes. Methods Mol Biol 1757, 399–470. 10.1007/978-1-4939-7737-6_14.

97. Huang, D.W., Sherman, B.T., Tan, Q., Collins, J.R., Alvord, W.G., Roayaei, J., Stephens, R., Baseler, M.W., Lane, H.C., and Lempicki, R.A. (2007). The DAVID Gene Functional Classification Tool: a novel biological module-centric algorithm to functionally analyze large gene lists. Genome Biol 8, R183. 10.1186/gb-2007-8-9-r183.

98. Huang, D.W., Sherman, B.T., Tan, Q., Kir, J., Liu, D., Bryant, D., Guo, Y., Stephens, R., Baseler, M.W., Lane, H.C., and Lempicki, R.A. (2007). DAVID Bioinformatics Resources: expanded annotation database and novel algorithms to better extract biology from large gene lists. Nucleic Acids Res 35, W169–175. 10.1093/nar/gkm415.

99. Sherman, B.T., Huang da, W., Tan, Q., Guo, Y., Bour, S., Liu, D., Stephens, R., Baseler, M.W., Lane, H.C., and Lempicki, R.A. (2007). DAVID Knowledgebase: a gene-centered database integrating heterogeneous gene annotation resources to facilitate high-throughput gene functional analysis. BMC Bioinformatics 8, 426. 10.1186/1471-2105-8-426.

100. Yang, W., Dierking, K., and Schulenburg, H. (2016). WormExp: a web-based application for a Caenorhabditis elegans-specific gene expression enrichment analysis. Bioinformatics 32, 943–945. 10.1093/bioinformatics/btv667.

101. Angeles-Albores, D., RY, N.L., Chan, J., and Sternberg, P.W. (2016). Tissue enrichment analysis for C. elegans genomics. BMC Bioinformatics 17, 366. 10.1186/s12859-016-1229-9.

102. Huang da, W., Sherman, B.T., and Lempicki, R.A. (2009). Systematic and integrative analysis of large gene lists using DAVID bioinformatics resources. Nat Protoc 4, 44–57. 10.1038/nprot.2008.211.

103. Huang da, W., Sherman, B.T., Zheng, X., Yang, J., Imamichi, T., Stephens, R., and Lempicki, R.A. (2009). Extracting biological meaning from large gene lists with DAVID. Curr Protoc Bioinformatics Chapter 13, Unit 13 11. 10.1002/0471250953.bi1311s27.

104. Schindelman, G., Fernandes, J.S., Bastiani, C.A., Yook, K., and Sternberg, P.W. (2011). Worm Phenotype Ontology: integrating phenotype data within and beyond the C. elegans community. BMC Bioinformatics 12, 32. 10.1186/1471-2105-12-32.

105. Singh, V., and Aballay, A. (2012). Endoplasmic Reticulum Stress Pathway Required for Immune Homeostasis Is Neurally Controlled by Arrestin-1. Journal of Biological Chemistry 287, 33191–33197. 10.1074/jbc.M112.398362.

106. Livak, K.J., and Schmittgen, T.D. (2001). Analysis of relative gene expression data using real-time quantitative PCR and the 2(-Delta Delta C(T)) Method. Methods 25, 402–408. 10.1006/meth.2001.1262.

107. Fay, D. (2006). Genetic mapping and manipulation: chapter 1--Introduction and basics. WormBook, 1-12. 10.1895/wormbook.1.90.1.

108. Fraser, A.G., Kamath, R.S., Zipperlen, P., Martinez-Campos, M., Sohrmann, M., and Ahringer, J. (2000). Functional genomic analysis of C. elegans chromosome I by systematic RNA interference. Nature 408, 325–330. 10.1038/35042517.

