## SupplementaryFigures for "Microbial odours activate protective immune response in *Caenorhabditis elegans* via specific olfactory neurons"

### Slide 1
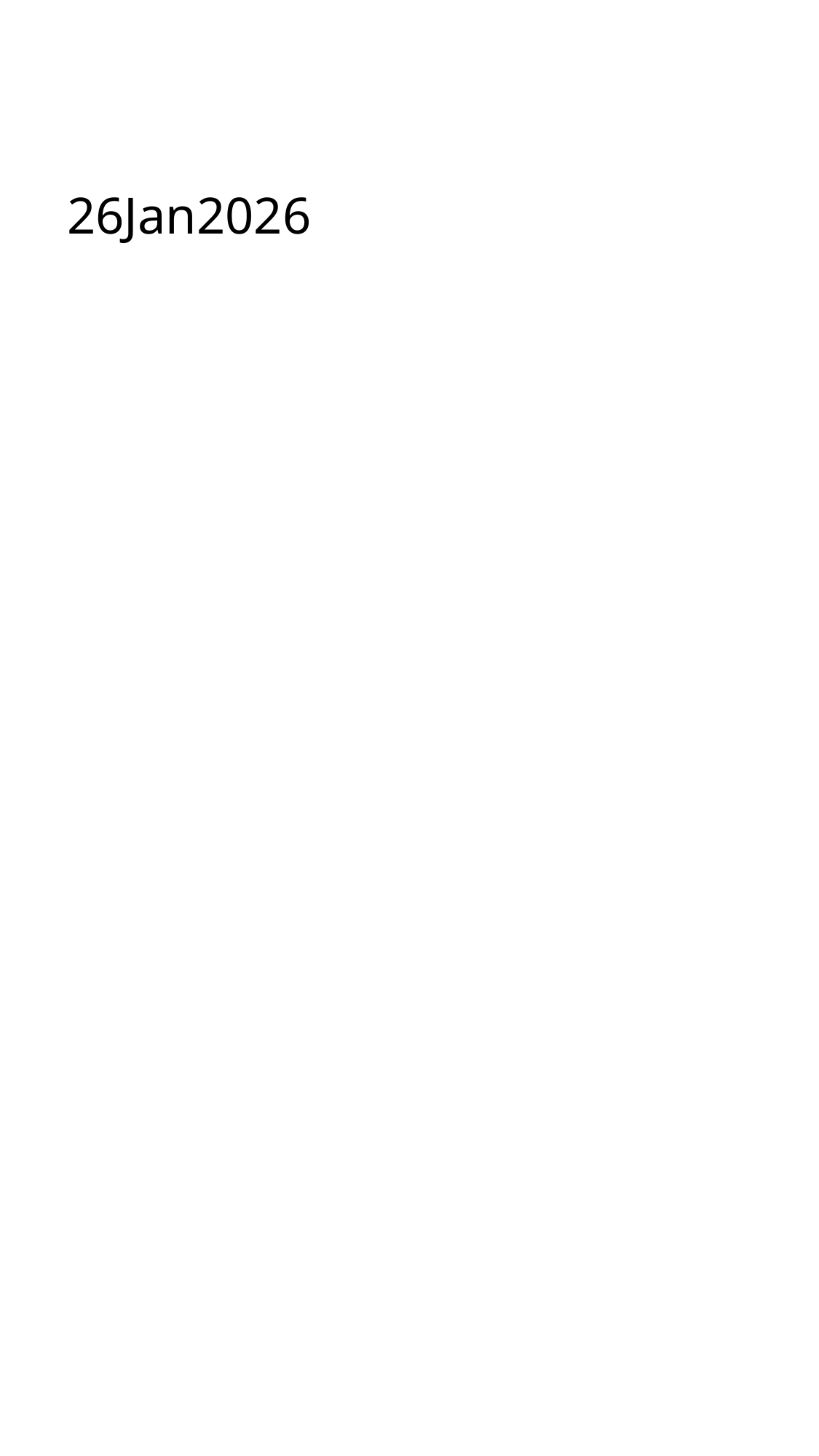

#### 26Jan2026

### Slide 2
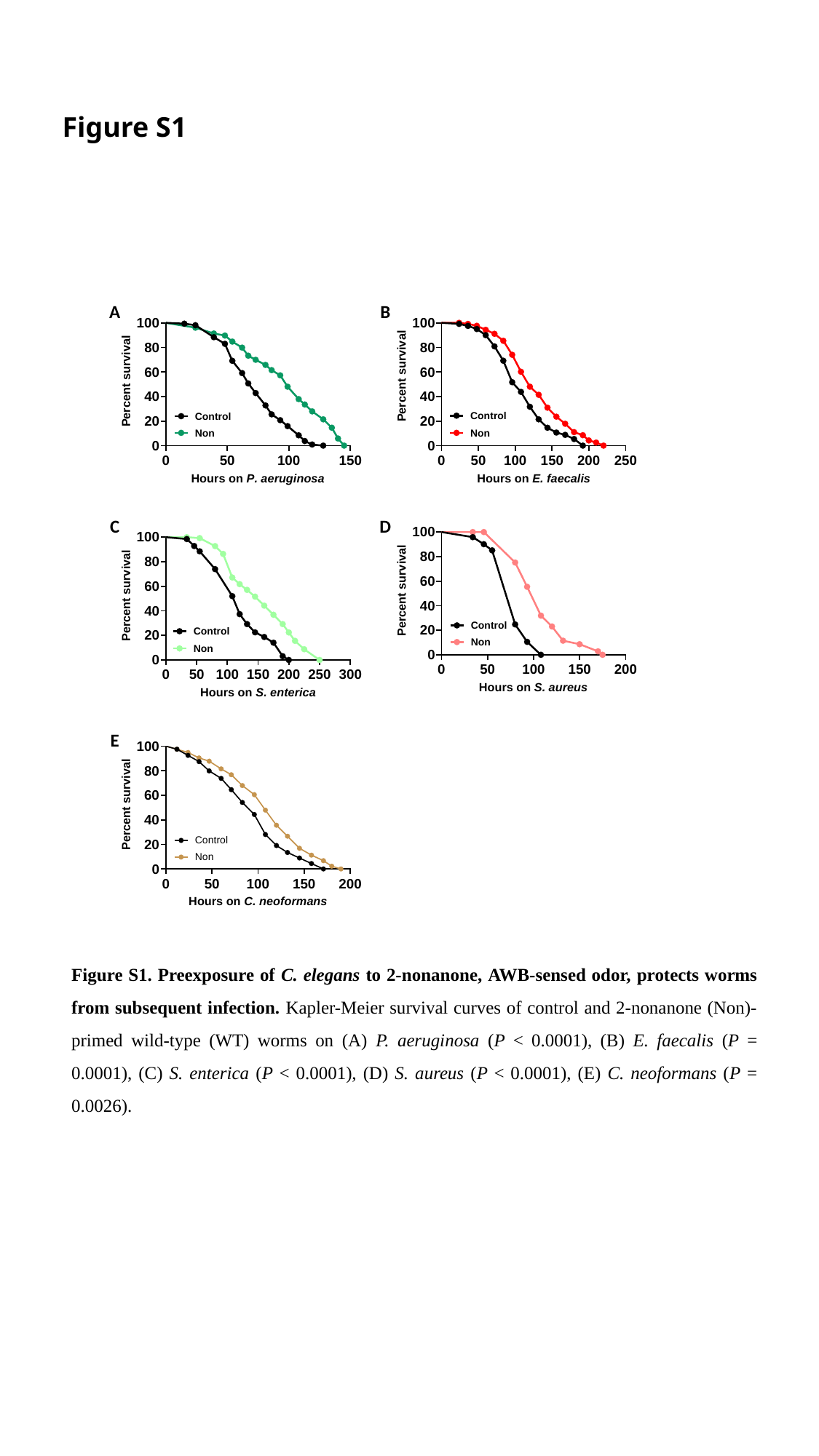

Figure S1
B
A
C
D
E
Figure S1. Preexposure of C. elegans to 2-nonanone, AWB-sensed odor, protects worms from subsequent infection. Kapler-Meier survival curves of control and 2-nonanone (Non)-primed wild-type (WT) worms on (A) P. aeruginosa (P < 0.0001), (B) E. faecalis (P = 0.0001), (C) S. enterica (P < 0.0001), (D) S. aureus (P < 0.0001), (E) C. neoformans (P = 0.0026).

### Slide 3
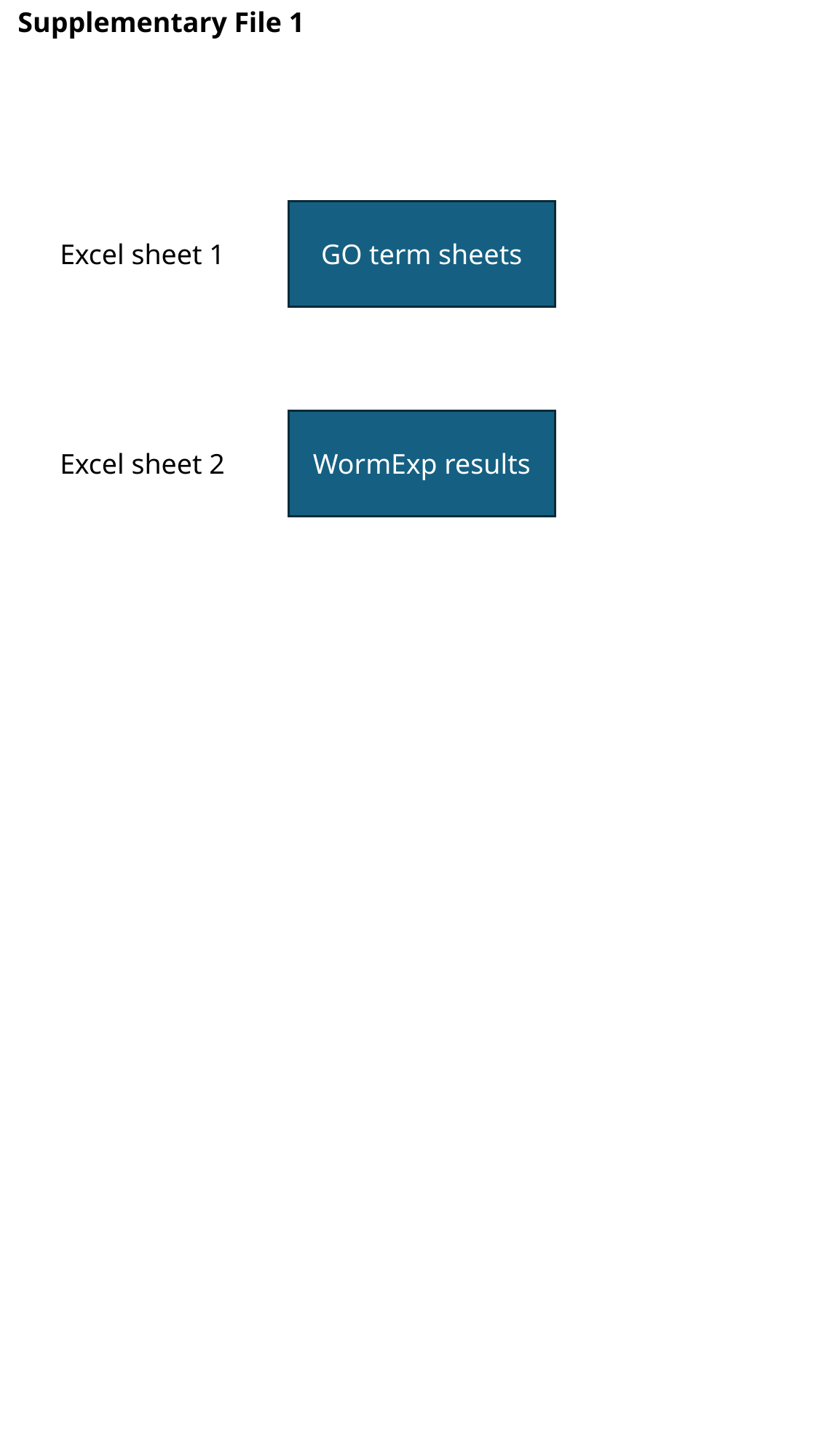

Supplementary File 1
GO term sheets
Excel sheet 1
WormExp results
Excel sheet 2

### Slide 4
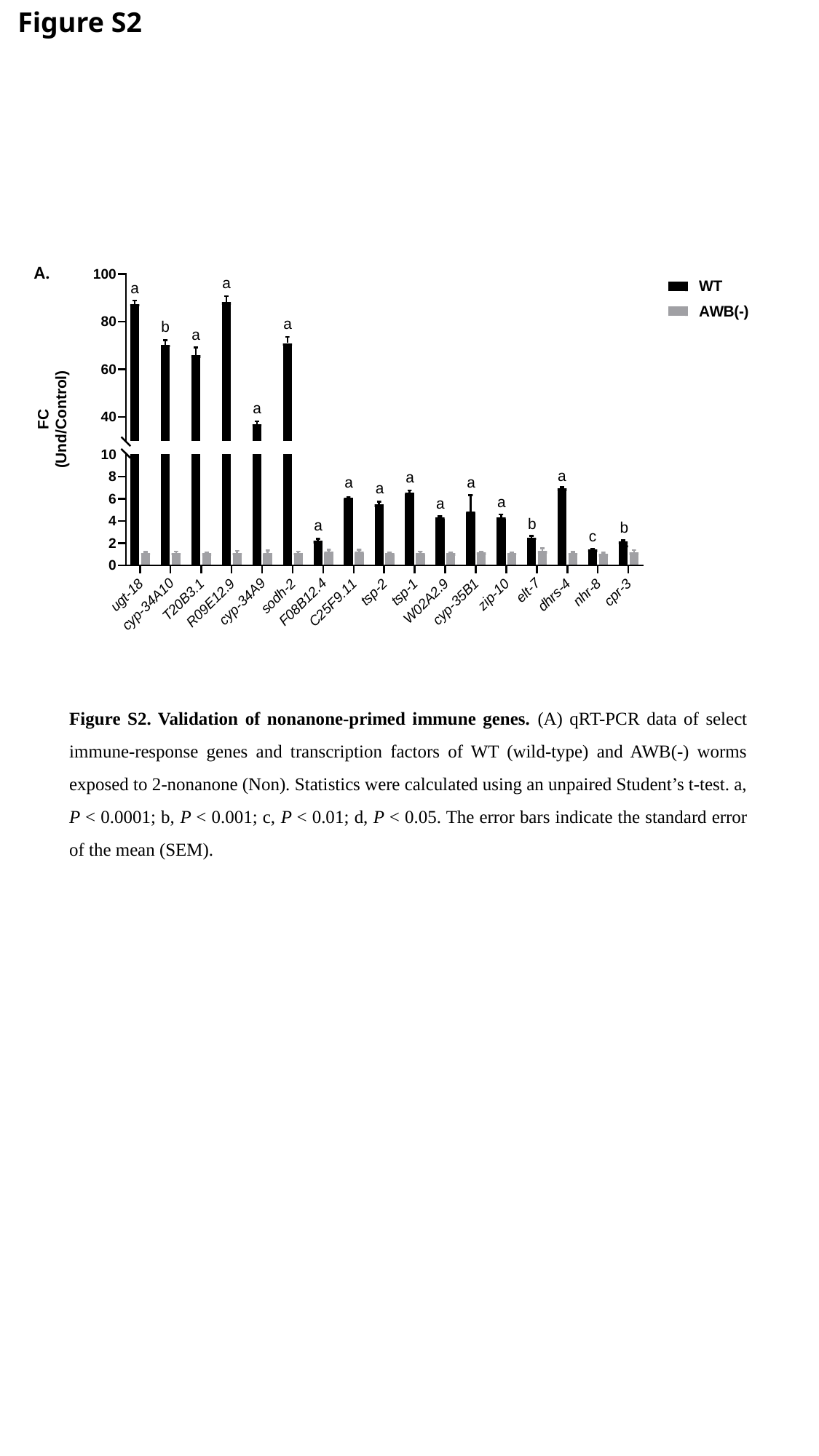

Figure S2
A.
Figure S2. Validation of nonanone-primed immune genes. (A) qRT-PCR data of select immune-response genes and transcription factors of WT (wild-type) and AWB(-) worms exposed to 2-nonanone (Non). Statistics were calculated using an unpaired Student’s t-test. a, P < 0.0001; b, P < 0.001; c, P < 0.01; d, P < 0.05. The error bars indicate the standard error of the mean (SEM).

### Slide 5
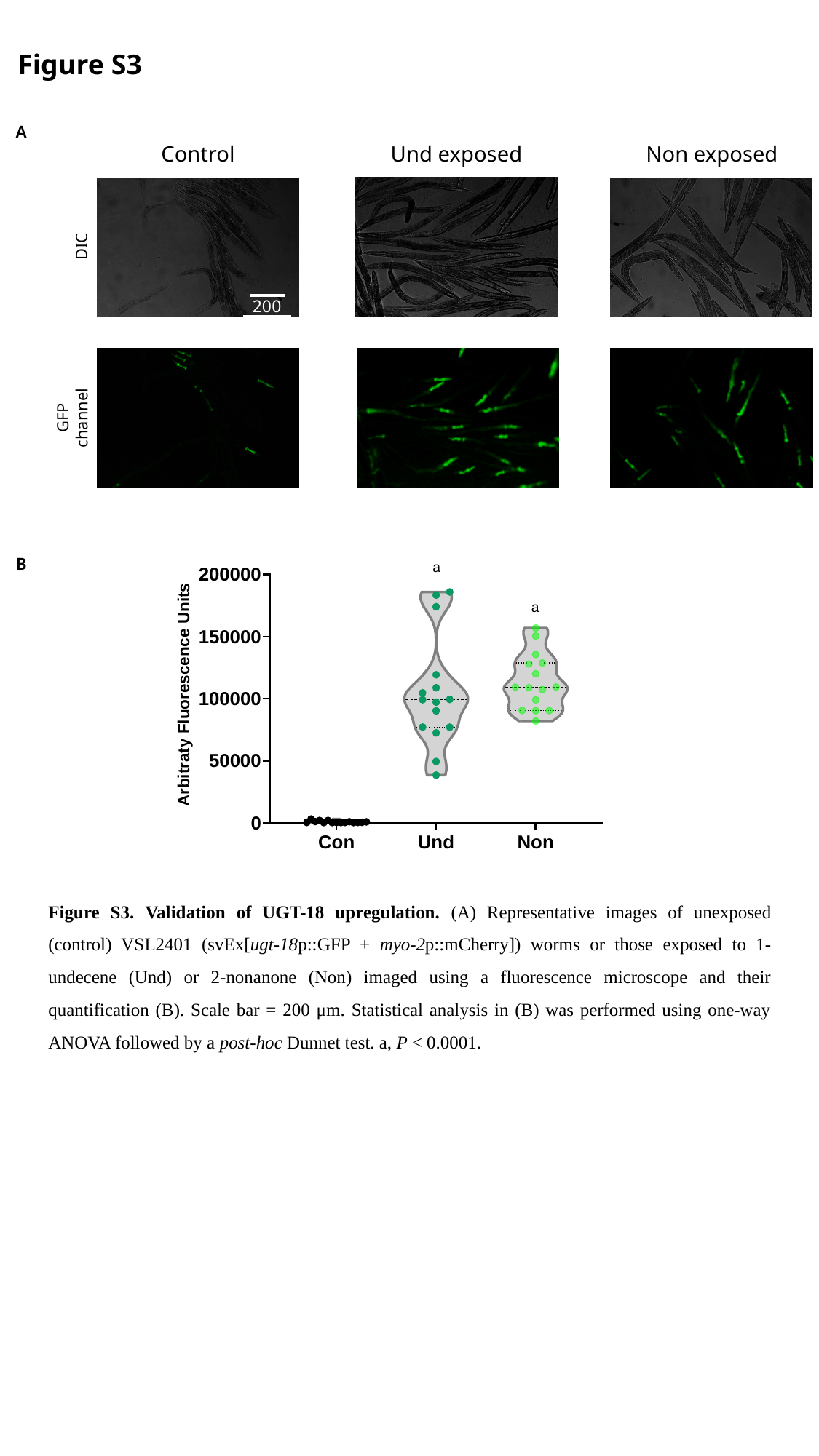

Figure S3
A
Control
Non exposed
Und exposed
DIC
200 μm
GFP
channel
B
Figure S3. Validation of UGT-18 upregulation. (A) Representative images of unexposed (control) VSL2401 (svEx[ugt-18p::GFP + myo-2p::mCherry]) worms or those exposed to 1-undecene (Und) or 2-nonanone (Non) imaged using a fluorescence microscope and their quantification (B). Scale bar = 200 μm. Statistical analysis in (B) was performed using one-way ANOVA followed by a post-hoc Dunnet test. a, P < 0.0001.

### Slide 6
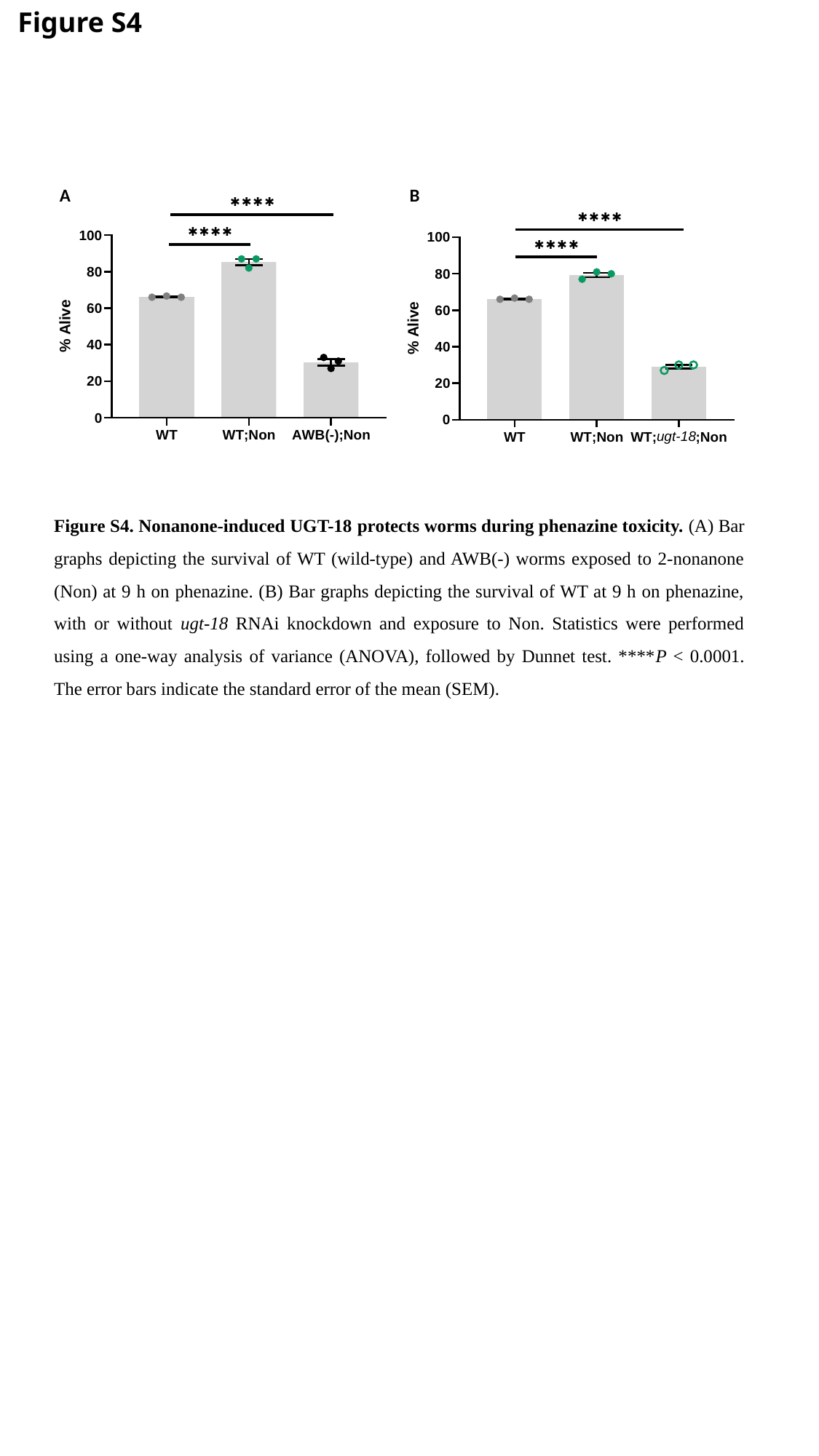

Figure S4
A
B
Figure S4. Nonanone-induced UGT-18 protects worms during phenazine toxicity. (A) Bar graphs depicting the survival of WT (wild-type) and AWB(-) worms exposed to 2-nonanone (Non) at 9 h on phenazine. (B) Bar graphs depicting the survival of WT at 9 h on phenazine, with or without ugt-18 RNAi knockdown and exposure to Non. Statistics were performed using a one-way analysis of variance (ANOVA), followed by Dunnet test. ****P < 0.0001. The error bars indicate the standard error of the mean (SEM).

### Slide 7
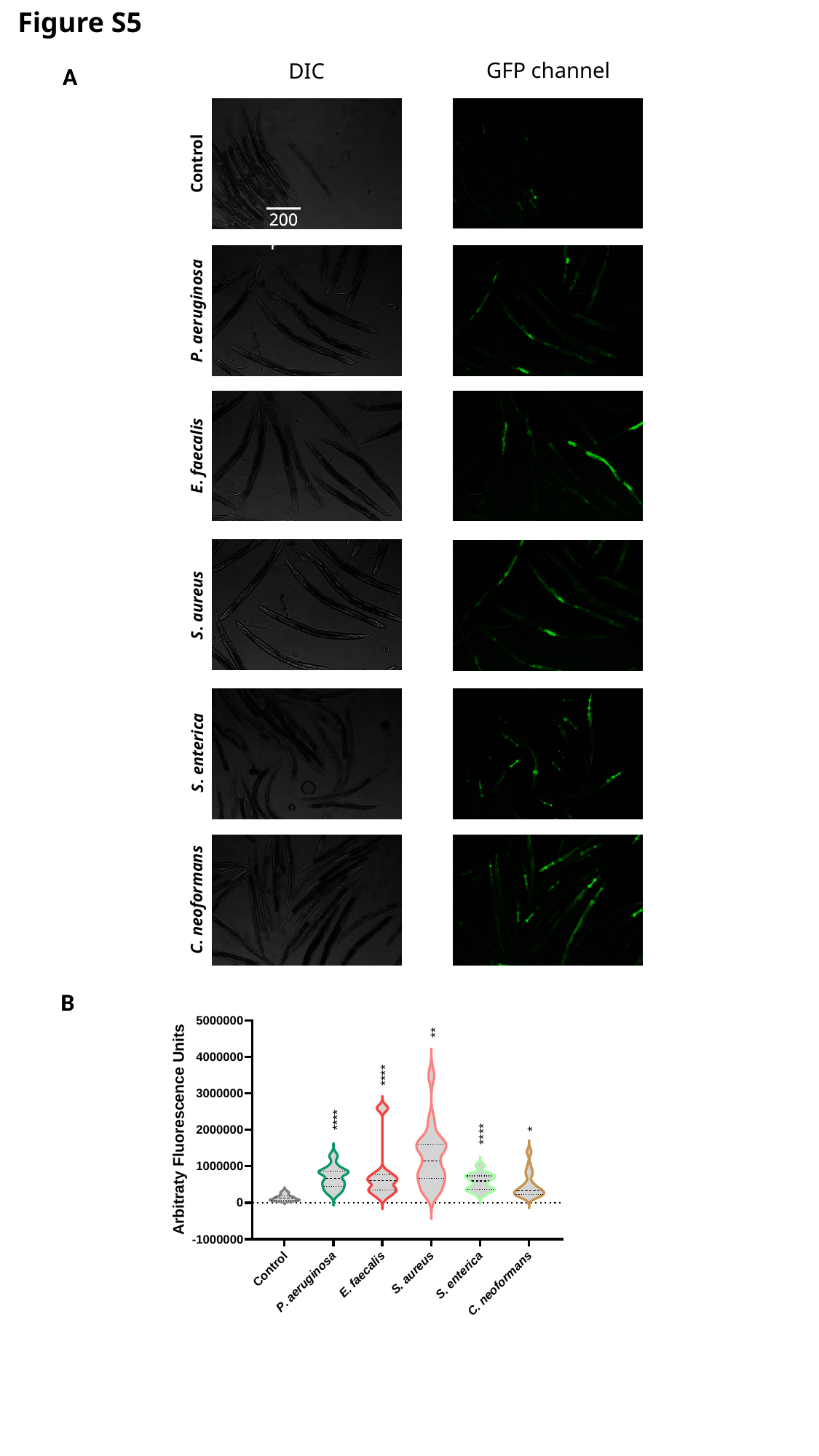

Figure S5
GFP channel
DIC
Control
P. aeruginosa
E. faecalis
S. aureus
S. enterica
C. neoformans
A
B
200 μm
A
200 μm
B

### Slide 8
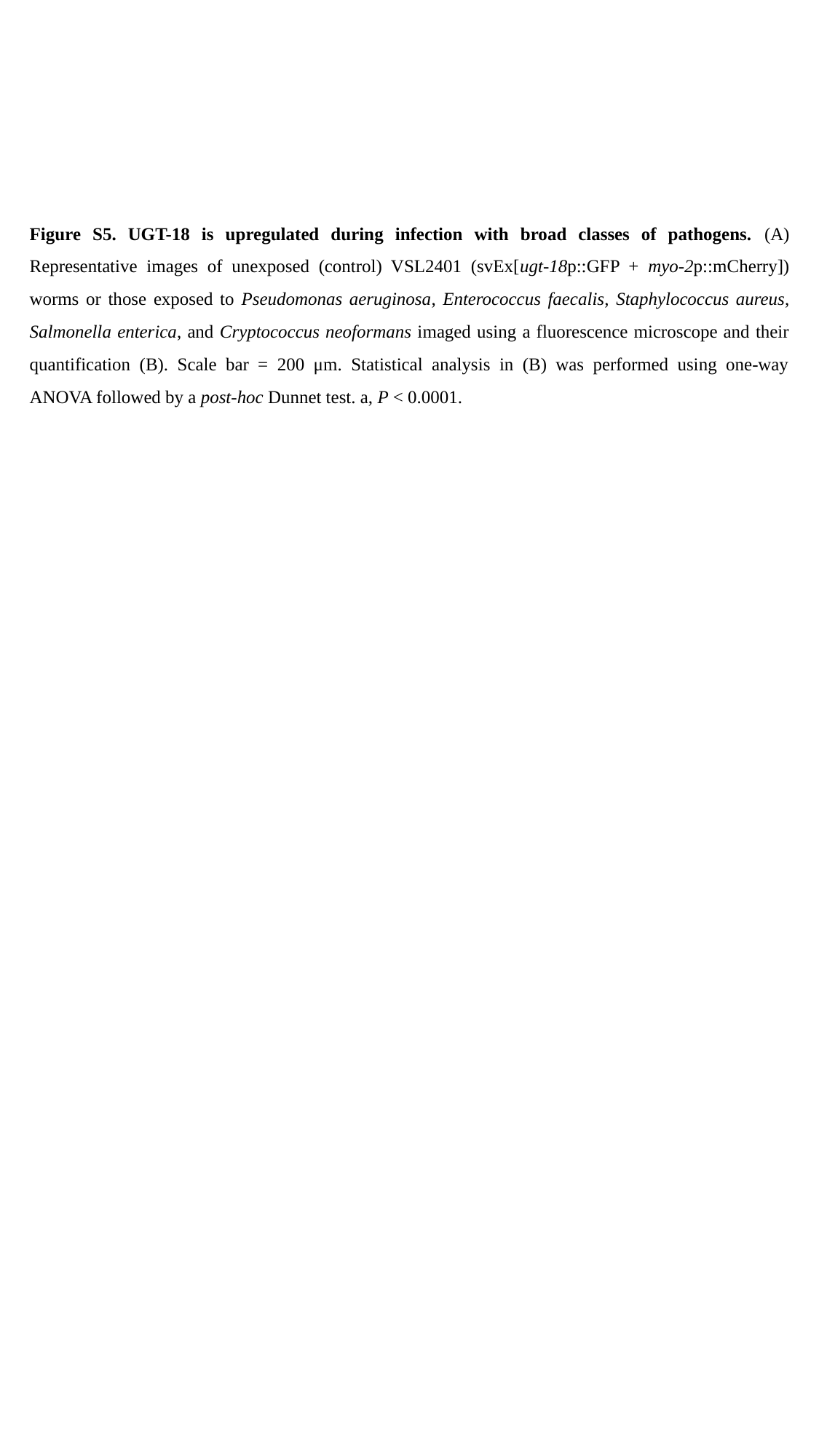

Figure S5. UGT-18 is upregulated during infection with broad classes of pathogens. (A) Representative images of unexposed (control) VSL2401 (svEx[ugt-18p::GFP + myo-2p::mCherry]) worms or those exposed to Pseudomonas aeruginosa, Enterococcus faecalis, Staphylococcus aureus, Salmonella enterica, and Cryptococcus neoformans imaged using a fluorescence microscope and their quantification (B). Scale bar = 200 μm. Statistical analysis in (B) was performed using one-way ANOVA followed by a post-hoc Dunnet test. a, P < 0.0001.
