## Supplementary Table S1 for "Microbial odours activate protective immune response in *Caenorhabditis elegans* via specific olfactory neurons"

**Table S1. Survival statistics**

| **Pathogen** | **Genotype/Condition** | **Number of animals (Assayed/Censored)** | **TD_50_ in Hours** | ***P*-value^#^** | **Figures** |
| --- | --- | --- | --- | --- | --- |
| *Pseudomonas aeruginosa* | N2/ Naïve  N2/ 1-Undecene | 111/14  115/12 | 70.01  94.01 | *P* < 0.0001 | **1B** |
|  | N2/ Naïve  N2/ 1-Undecene | 110/10  120/5 | 69.72  95.11 | *P* < 0.0001 |  |
|  | N2/ Naïve  N2/ 1-Undecene | 108/10  111/13 | 72.82  97.21 | *P* < 0.0001 |  |
| *P*. *aeruginosa* | AWB(-)/ Naïve  AWB(-)/ 1-Undecene | 111/4  116/4 | 74.33  76.56 | *P* = 0.1712 | **1C** |
|  | AWB(-)/ Naïve  AWB(-)/ 1-Undecene | 107/23  111/8 | 69.46  71.23 | *P* = 0.2644 |  |
|  | AWB(-)/ Naïve  AWB(-)/ 1-Undecene | 90/23  103/13 | 70.45  71.23 | *P* = 0.6923 |  |
| *Enterococcus faecalis* | N2/ Naïve  N2/ 1-Undecene | 111/10  114/12 | 98.61  140.46 | *P* < 0.0001 | **1D** |
|  | N2/ Naïve  N2/ 1-Undecene | 108/10  120/4 | 99.12  142.63 | *P* < 0.0001 |  |
|  | N2/ Naïve  N2/ 1-Undecene | 110/20  120/7 | 95.23  144.21 | *P* < 0.0001 |  |
| *Salmonella enterica*  *S*. *enterica* | N2/ Naïve  N2/ 1-Undecene | 130/7  106/20 | 97.92  164.12 | *P* < 0.0001 | **1E** |
|  | N2/ Naïve  N2/ 1-Undecene | 120/1  109/10 | 99.19  163.27 | *P* < 0.0001 |  |
|  | N2/ Naïve  N2/ 1-Undecene | 111/8  120/6 | 101.42  154.26 | *P* < 0.0001 |  |
| *Staphylococcus aureus* | N2/ Naïve  N2/ 1-Undecene | 113/24  107/23 | 60.33  85.33 | *P* < 0.0001 | **1F** |
|  | N2/ Naïve  N2/ 1-Undecene | 120/5  117/10 | 64.46  86.12 | *P* < 0.0001 |  |
|  | N2/ Naïve  N2/ 1-Undecene | 120/10  98/28 | 63.85  88.35 | *P* < 0.0001 |  |
| *Cryptococcus neoformans* | N2/ Naïve  N2/ 1-Undecene | 99/23  95/29 | 88.67  111.44 | *P* = 0.0002 | **1E** |
|  | N2/ Naïve  N2/ 1-Undecene | 101/24  113/12 | 90.63  113.42 | *P* < 0.0001 |  |
|  | N2/ Naïve  N2/ 1-Undecene | 102/32  121/7 | 92.34  119.12 | *P* < 0.0001 |  |
| *P*. *aeruginosa* | N2/ Naïve  N2/ 2-Nonanone | 127/35  115/13 | 67.17  97.75 | *P* < 0.0001 | **S1A** |
|  | N2/ Naïve  N2/ 2-Nonanone | 120/4  116/6 | 69.25  92.35 | *P* < 0.0001 |  |
|  | N2/ Naïve  N2/ 2-Nonanone | 118/4  96/19 | 70.13  98.46 | *P* < 0.0001 |  |
| *E*. *faecalis*  *E*. *faecalis* | N2/ Naïve  N2/ 2-Nonanone | 111/10  122/4 | 98.33  117.64 | *P* < 0.0001 | **S1B** |
|  | N2/ Naïve  N2/ 2-Nonanone | 120/0  124/4 | 97.25  120.45 | *P* < 0.0001 |  |
|  | N2/ Naïve  N2/ 2-Nonanone | 121/4  103/15 | 92.46  117.35 | *P* < 0.0001 |  |
| *S*. *enterica* | N2/ Naïve  N2/ 2-Nonanone | 134/6  113/33 | 109.83  148.50 | *P* < 0.0001 | **S1C** |
|  | N2/ Naïve  N2/ 2-Nonanone | 121/3  117/20 | 111.06  147.24 | *P* < 0.0001 |  |
|  | N2/ Naïve  N2/ 2-Nonanone | 102/12  108/15 | 112.34  150.21 | *P* < 0.0001 |  |
| *S*. *aureus* | N2/ Naïve  N2/ 2-Nonanone | 119/3  96/29 | 69.78  96.33 | *P* < 0.0001 | **S1D** |
|  | N2/ Naïve  N2/ 2-Nonanone | 118/13  105/14 | 72.45  99.34 | *P* < 0.0001 |  |
|  | N2/ Naïve  N2/ 2-Nonanone | 106/23  111/10 | 64.26  92.35 | *P* < 0.0001 |  |
| *C*. *neoformans* | N2/ Naïve  N2/ 2-Nonanone | 99/23  86/34 | 88.67  106.22 | *P* = 0.0026 | **S1E** |
|  | N2/ Naïve  N2/ 2-Nonanone | 101/24  120/4 | 90.63 | *P* < 0.0001 |  |
|  | N2/ Naïve  N2/ 2-Nonanone | 102/32  118/12 | 92.34 | *P* < 0.001 |  |
| *P*. *aeruginosa*  *P*. *aeruginosa* | N2/ Naïve  N2/ Diacetyl  CX3877/ Naïve  CX3877/ Diacetyl | 94/19  117/9  127/2  95/0 | 67.33  69.56  67.41  80.44 | *P* < 0.0001 | **2B** |
|  | N2/ Naïve  N2/ Diacetyl  CX3877/ Naïve  CX3877/ Diacetyl | 111/14  127/35  108/0  120/1 | 70.47  67.38  66.83  104.83 | *P* < 0.0001 |  |
|  | N2/ Naïve  N2/ Diacetyl  CX3877/ Naïve  CX3877/ Diacetyl | 94/19  104/11  65/0  62/3 | 57.25  59.12  58.08  96.33 | *P* < 0.0001 |  |
| *E*. *faecalis* | N2/ Naïve  N2/ Diacetyl  CX3877/ Naïve  CX3877/ Diacetyl | 111/12  106/20  100/15  96/29 | 69.58  67.22  67.36  65.69 | *P* < 0.0001 | **2C** |
|  | N2/ Naïve  N2/ Diacetyl  CX3877/ Naïve  CX3877/ Diacetyl | 111/10  130/5  127/1  115/10 | 98.33  99.44  99.23  126.83 | *P* < 0.0001 |  |
|  | N2/ Naïve  N2/ Diacetyl  CX3877/ Naïve  CX3877/ Diacetyl | 121/12  108/20  109/11  112/12 | 98.91  98.10  97.23  125.46 | *P* < 0.0001 |  |
| *S*. *enterica*  *S*. *enterica* | N2/ Naïve  N2/ Diacetyl  CX3877/ Naïve  CX3877/ Diacetyl | 130/7  93/29  114/6  79/2 | 98.33  96.67  107.64  167.68 | *P* < 0.0001 | **2D** |
|  | N2/ Naïve  N2/ Diacetyl  CX3877/ Naïve  CX3877/ Diacetyl | 122/10  124/7  120/9  127/1 | 108.18  106.43  102.23  157.73 | *P* < 0.0001 |  |
|  | N2/ Naïve  N2/ Diacetyl  CX3877/ Naïve  CX3877/ Diacetyl | 106/12  118/11  111/10  117/19 | 103.56  104.35  99.38  148.73 | *P* < 0.0001 |  |
| *S*. *aureus* | N2/ Naïve  N2/ Diacetyl  CX3877/ Naïve  CX3877/ Diacetyl | 90/37  76/49  108/4  118/22 | 53.50  53.50  54.33  76.17 | *P* < 0.0001 | **2E** |
|  | N2/ Naïve  N2/ Diacetyl  CX3877/ Naïve  CX3877/ Diacetyl | 120/10  108/21  98/23  80/35 | 58.19  58.92  55.69  77.23 | *P* < 0.0001 |  |
|  | N2/ Naïve  N2/ Diacetyl  CX3877/ Naïve  CX3877/ Diacetyl | 108/22  110/27  89/20  100/15 | 55.82  56.127  54.24  80.12 | *P* < 0.0001 |  |
| *P*. *aeruginosa*  *P*. *aeruginosa* | AWB::HisCl/ No HA  AWB::HisCl/ 0-24 h HA  AWB::HisCl/ 24-48 h HA | 112/12  112/12  68/33 | 74.58  35.56  70.833 | *P* < 0.0001 | **3E** |
|  | AWB::HisCl/ No HA  AWB::HisCl/ 0-24 h HA  AWB::HisCl/ 24-48 h HA | 142/0  118/13  107/13 | 70.33  55.42  71.98 | *P* < 0.0001 |  |
|  | AWB::HisCl/ No HA  AWB::HisCl/ 0-24 h HA  AWB::HisCl/ 24-48 h HA | 103/12  119/4  122/5 | 66.72  45.25  64.81 | *P* < 0.0001 |  |
| *P*. *aeruginosa* | AWB::HisCl/ No HA  AWB::HisCl/ 0-8 HA  AWB::HisCl/ 8-24 HA  AWB::HisCl/ 0-24 HA | 97/15  96/7  98/12  92/5 | 61.11  44.65  47.15  35.49 | *P* < 0.0001 | **3F** |
|  | AWB::HisCl/ No HA  AWB::HisCl/ 0-8 HA  AWB::HisCl/ 8-24 HA  AWB::HisCl/ 0-24 HA | 142/0  127/9  145/3  118/13 | 70.33  59.08  63.08  55.42 | *P* < 0.0001 |  |
|  | AWB::HisCl/ No HA  AWB::HisCl/ 0-8 HA  AWB::HisCl/ 8-24 HA  AWB::HisCl/ 0-24 HA | 97/14  116/0  120/5  119/4 | 69.67  51.17  50.69  45.25 | *P* < 0.0001 |  |
| *P*. *aeruginosa* | N2/ EV  N2/ *ugt-18* | 92/28  124/1 | 64.84  14.12 | *P* < 0.0001 | **6A** |
|  | N2/ EV  N2/ *ugt-18* | 93/11  120/0 | 63.67  12.58 | *P* < 0.0001 |  |
|  | N2/ EV  N2/ *ugt-18* | 92/28  124/1 | 64.41  14.30 | *P* < 0.0001 |  |
| *P*. *aeruginosa*  *P*. *aeruginosa* | N2/ EV  N2/ *R09E12.9* | 103/27  100/20 | 71.42  63.17 | *P* < 0.0001 | **6B** |
|  | N2/ EV  N2/ *R09E12.9* | 110/21  106/12 | 68.12  51.28 | *P* < 0.0001 |  |
|  | N2/ EV  N2/ *R09E12.9* | 97/23  88/25 | 65.27  50.11 | *P* < 0.0001 |  |
| *P*. *aeruginosa* | N2/ EV  N2/ *sodh-2* | 92/28  98/22 | 64.23  57.75 | *P* = 0.0015 | **6C** |
|  | N2/ EV  N2/ *sodh-2* | 106/14  107/18 | 79.75  73.17 | *P* < 0.001 |  |
|  | N2/ EV  N2/ *sodh-2* | 93/11  115/30 | 63.67  52.47 | *P* < 0.001 |  |
| *P*. *aeruginosa* | N2/ EV  N2/ *T20B3.1* | 100/20  114/9 | 71  53.08 | *P* < 0.0001 | **6D** |
|  | N2/ EV  N2/ *T20B3.1* | 87/29  91/24 | 68.53  57.23 | *P* < 0.0001 |  |
|  | N2/ EV  N2/ *T20B3.1* | 120/2  111/15 | 72.35  55.64 | *P* < 0.0001 |  |
| *P*. *aeruginosa* | N2/ EV  N2/ *W02A2.9* | 87/33  91/31 | 81.88  66.54 | *P* = 0.0030 | **6E** |
|  | N2/ EV  N2/ *W02A2.9* | 99/20  91/24 | 70.53  62.15 | *P* < 0.0001 |  |
|  | N2/ EV  N2/ *W02A2.9* | 109/12  110/27 | 71.78  60.73 | *P* < 0.001 |  |
| *P*. *aeruginosa*  *P*. *aeruginosa* | N2/ EV  N2/ *cpr-3* | 87/33  96/24 | 86.75  67.25 | *P* < 0.0001 | **6F** |
|  | N2/ EV  N2/ *cpr-3* | 106/12  110/7 | 80.62  62.53 | *P* < 0.0001 |  |
|  | N2/ EV  N2/ *cpr-3* | 97/21  89/25 | 72.54  53.74 | *P* < 0.0001 |  |
| *P*. *aeruginosa* | N2/ EV  N2/ *tsp-1* | 109/1  116/0 | 54.33  51.06 | *P* = 0.1348 | **6G** |
|  | N2/ EV  N2/ *tsp-1* | 120/4  126/1 | 55.72  52.53 | *P* = 0.2763 |  |
|  | N2/ EV  N2/ *tsp-1* | 110/27  90/38 | 54.72  53.98 | *P* = 0.5672 |  |
| *P*. *aeruginosa* | N2/ EV  N2/ *F08B12.4* | 89/36  89/36 | 78.71  72.66 | *P* = 0.4263 | **6H** |
|  | N2/ EV  N2/ *F08B12.4* | 120/2  109/21 | 54.82  53.83 | *P* = 0.2612 |  |
|  | N2/ EV  N2/ *F08B12.4* | 102/18  111/9 | 58.82  59.98 | *P* = 0.6739 |  |
| *P*. *aeruginosa* | N2/ EV  N2/ *C25F9.11* | 111/1  105/11 | 55.94  57.72 | *P* = 0.0614 | **6I** |
|  | N2/ EV  N2/ *C25F9.11* | 102/8  110/11 | 57.90  55.67 | *P* = 0.1621 |  |
|  | N2/ EV  N2/ *C25F9.11* | 120/11  99/31 | 71.27  72.92 | *P* = 0.0798 |  |
| Phenazine | N2/ DMSO  N2/ Phenazine | 60/0  60/0 | NA  10.58 | *P* < 0.0001 | **7A** |
|  | N2/ DMSO  N2/ Phenazine | 60/0  60/0 | NA  10.71 | *P* < 0.0001 |  |
|  | N2/ DMSO  N2/ Phenazine | 60/0  60/0 | NA  10.92 | *P* < 0.0001 |  |
| Phenazine | N2/ EV  N2/ *ugt-18*  AWB(-)/ EV  AWB(-)/ *ugt-18* | 60/0  60/0  60/0  60/0 | 10.44  7.31  6.43  5.64 | *P* < 0.0001 | **7C** |
|  | N2/ EV  N2/ *ugt-18*  AWB(-)/ EV  AWB(-)/ *ugt-18* | 60/0  60/0  60/0  60/0 | 10.76  7.91  7.21  6.93 | *P* < 0.0001 |  |
|  | N2/ EV  N2/ *ugt-18*  AWB(-)/ EV  AWB(-)/ *ugt-18* | 60/0  60/0  60/0  60/0 | 10.11  7.12  6.90  6.31 | *P* < 0.0001 |  |
| Phenazine | N2  N2/ Und  N2/ *ugt-18*/ Und | 60/0  60/0  60/0 | 10.24  14.933  7.80 | *P* < 0.0001 | **7F** |
|  | N2  N2/ Und  N2/ *ugt-18*/ Und | 60/0  60/0  60/0 | 10.69  15.82  7.23 | *P* < 0.0001 |  |
|  | N2  N2/ Und  N2/ *ugt-18*/ Und | 60/0  60/0  60/0 | 10.46  14.46  7.47 | *P* < 0.0001 |  |
| *E*. *faecalis*  *E*. *faecalis* | N2/ EV  N2/ *ugt-18* | 108/22  95/20 | 65.21  48.47 | *P* < 0.0001 | **8A** |
|  | N2/ EV  N2/ *ugt-18* | 70/25  69/2 | 76.08  59.08 | *P* = 0.0001 |  |
|  | N2/ EV  N2/ *ugt-18* | 112/12  108/21 | 74.5  54.83 | *P* < 0.0001 |  |
| *S*. *enterica* | N2/ EV  N2/ *ugt-18* | 99/23  86/34 | 105.89  88.89 | *P* < 0.0026 | **8B** |
|  | N2/ EV  N2/ *ugt-18* | 94/26  95/28 | 95  83.11 | *P* = 0.0430 |  |
|  | N2/ EV  N2/ *ugt-18* | 101/29  84/51 | 137.87  96.43 | *P* < 0.0001 |  |
| *S*. *aureus* | N2/ EV  N2/ *ugt-18* | 67/33  84/26 | 82.67  68.58 | *P* = 0.0002 | **8C** |
|  | N2/ EV  N2/ *ugt-18* | 132/43  126/21 | 72.25  61.08 | *P* < 0.0001 |  |
|  | N2/ EV  N2/ *ugt-18* | 112/12  85/5 | 74.42  59.25 | *P* < 0.0001 |  |
| *C*. *neoformans* | N2/ EV  N2/ *ugt-18* | 94/26  118/12 | 84.56  65.22 | *P* = 0.0002 | **8D** |
|  | N2/ EV  N2/ *ugt-18* | 109/11  108/12 | 88.36  69.21 | *P* < 0.001 |  |
|  | N2/ EV  N2/ *ugt-18* | 120/2  111/6 | 87.1  64.25 | *P* < 0.0001 |  |

NA, Not applicable

^#^Some of the analysis were performed with the same control data as the experiments were performed together.
