## Supplementary Table S3 for "Microbial odours activate protective immune response in *Caenorhabditis elegans* via specific olfactory neurons"

**Table S3. List of *C*. *elegans* strains used in this study**

| **Strain Name** | **Genotype** | **Reference** |
| --- | --- | --- |
| N2 / WT | N2 | **S. Brenner, 1974** |
| JN1715/AWB(-) | peIs1715 [*str-1p*::mCasp-1 + *unc-122p*::GFP] | **(Yoshida et al., 2012)** |
| PS1799 | syIs666 [*str-1p*::NLS::GAL4(sk)::VP64::*let-858* 3'UTR + *unc-122p*::RFP + 1kb DNA ladder (NEB)] + *unc-122p*::GFP + 1kb DNA ladder(NEB)] | **(H. Wang et al., 2017)** |
| PS8565 | syIs666 [*str-1p*::NLS::GAL4(sk)::VP64::*let-858* 3'UTR + *unc-122p*::RFP + 1kb DNA ladder (NEB)] | **(H. Wang et al., 2017)** |
| VSL2399/  AWB::HisCl | peIs1715 [*str-1p*::mCasp-1 + *unc-122p*::GFP] + syIs371 [15xUAS::Δpes-10::HisCl1::SL2::GFP::let-858 3'UTR + *unc-122p*::GFP + 1kb DNA ladder(NEB)] | **This study** |
| CX3877/ AWBp::ODR-10 | kyIs156 [*str-1p*::*odr-10*(cDNA)::GFP] | **(Maniar et al., 2011)** |
| VSL2403 | svEx [*ugt-18p*::GFP + *myo-2p*::mCherry] | **This study** |
